# Diversification of mammalian deltaviruses by host shifting

**DOI:** 10.1101/2020.06.17.156745

**Authors:** Laura M. Bergner, Richard J. Orton, Alice Broos, Carlos Tello, Daniel J. Becker, Jorge E. Carrera, Arvind H. Patel, Roman Biek, Daniel G. Streicker

## Abstract

Hepatitis delta virus (HDV) is an unusual RNA agent that replicates using host machinery but exploits hepatitis B virus (HBV) to mobilize its spread within and between hosts. In doing so, HDV enhances the virulence of HBV. How this seemingly improbable hyper-parasitic lifestyle emerged is unknown, but underpins the likelihood that HDV and related deltaviruses may alter other host-virus interactions. Here, we show that deltaviruses diversify by transmitting between mammalian species. Among 96,695 RNA sequence datasets, deltaviruses infected bats, rodents and an artiodactyl from the Americas, but were absent from geographically overrepresented Old World representatives of each mammalian order, suggesting a relatively recent diversification within the Americas. Consistent with diversification by host shifting, both bat and rodent-infecting deltaviruses were paraphyletic and co-evolutionary modeling rejected co-speciation with mammalian hosts. In addition, a two-year field study showed common vampire bats in Peru were infected by two divergent deltaviruses, indicating multiple introductions to a single host species. One vampire bat-associated deltavirus was detected in the saliva of up to 35% of individuals, formed phylogeographically compartmentalized clades, and infected a sympatric bat, illustrating horizontal transmission within and between species on ecological timescales. Consistent absence of HBV-like viruses in two deltavirus-infected bat species indicated acquisitions of novel viral associations during the divergence of bat and human-infecting deltaviruses. Our analyses support an American zoonotic origin of HDV and reveal prospects for future cross-species emergence of deltaviruses. Given their peculiar life history, deltavirus host shifts will have different constraints and disease outcomes compared to ordinary animal pathogens.

**Significance Statement:** Satellites are virus-like agents which require both a host and a virus to complete their life cycle. The only human-infecting satellite is hepatitis delta virus (HDV), which exacerbates liver disease in patients co-infected with hepatitis B virus (HBV). How HDV originated is a longstanding evolutionary puzzle. Using terabase-scale data mining, co-evolutionary analyses, and field studies in bats, we show that deltaviruses can jump between highly divergent host species. Our results further suggest that the contemporary association between HDV and HBV likely arose following zoonotic transmission from a yet undiscovered animal reservoir in the Americas. Plastic host and virus associations open prospects that deltaviruses might alter the virulence of multiple viruses in multiple host species.

## Introduction

Hepatitis delta virus (HDV), the only member of the only species (*Hepatitis delta virus*) in the genus *Deltavirus*, is a globally-distributed human pathogen which causes the most severe form of viral hepatitis in an estimated 20 million people (1). Unlike typical viruses, HDV is an obligate ‘satellite’ virus that is replicated by diverse host cells, but requires the envelope of an unrelated ‘helper’ virus (classically hepatitis B virus, HBV, family *Hepadnaviridae*) for cellular entry, egress and transmission (1). The peculiar life history of HDV together with its lack of sequence homology to known viral groups have made the evolutionary origins of HDV a longstanding puzzle. Geographic associations of most HDV genotypes point to an Old World origin. Yet, historical explanations of the mechanistic origin of HDV spanned from emergence from the mRNA of a HBV-infected human (2) to ancient evolution from viroids (circular, single-stranded RNA pathogens of plants) (3). More recently, discoveries of HDV-like genomes in vertebrates and invertebrates (4-7) overturned the decades-long belief that deltaviruses exclusively infect humans. These discoveries also suggested new models of deltavirus evolution, in which these satellites either co-speciated with their hosts over ancient timescales or possess an unrecognized capacity for host shifting which would imply their potential to emerge in novel species. The latter scenario has been presumed unlikely since either both satellite and helper would need to be compatible with the novel host, or deltaviruses would need to simultaneously switch host species and helper virus, possibly altering the virulence of newly acquired helpers as a result.

Efforts to distinguish competing evolutionary hypotheses for deltaviruses have been precluded by the remarkably sparse distribution of currently-known HDV-like agents across the animal tree of life. Single representatives are reported from arthropods (Subterranean termite, *Schedorhinotermes intermedius*), fish (a pooled sample from multiple species), birds (a pooled sample from 3 duck species, *Anas gracilis, A. castanea, A. superciliosa*), reptiles (Common boa, *Boa constrictor*) and mammals (Tome’s spiny rat, *Proechimys semispinosus*), and only two are known from amphibians (Asiatic toad, *Bufo gargarizans*; Chinese fire belly newt, *Cynops orientalis*) (4-7). Most share minimal homology with HDV, even at the protein level (< 25%), frustrating robust phylogenetic re-constructions of evolutionary histories (see SI Appendix, Fig. S1). On the one hand, the distribution of deltaviruses may reflect rare host shifting events among divergent taxa. Alternatively, reliance on untargeted metagenomic sequencing (a relatively new and selectively applied tool) to find novel species may mean that the distribution of deltaviruses in nature is largely incomplete (8, 9). Additional taxa could reveal ancient co-speciation of HDV-like agents with their hosts or evidence for host shifting.

## Results

We sought to fill gaps in the evolutionary history of mammalian deltaviruses, the group most likely to clarify the origins of HDV. We used a two-pronged approach (Materials and Methods). First, we used data from Serratus, a newly developed bioinformatic platform which screens RNA sequences from the NCBI Short Read Archive (SRA) for similarity to known viruses and which is described by Edgar *et al*. (10). We focused on search results from 96,695 transcriptomic and metagenomic datasets, comprising 348 terabases of RNA sequences from 403 species across 24 mammalian orders (22 terrestrial, 2 aquatic; see SI Appendix, Data S1). Although domesticated animals comprised the largest single fraction of the dataset (67.2%), remaining data were from a variety of globally-distributed species (Fig 1A,B). Our second search was prompted by our earlier detection of uncharacterized deltavirus-like sequences in a Neotropical bat (11) and evidence of under-representation in the volume of Neotropical bat data in the SRA (Fig. 1A). We therefore carried out metagenomic sequencing of 23 frugivorous, insectivorous, nectarivorous, and sanguivorous bat species from Peru, using 59 samples available within our laboratory (see SI Appendix, Table S1). All datasets containing sequences with significant protein homology to deltaviruses were subjected to *de novo* genome assembly.

**Figure 1.**
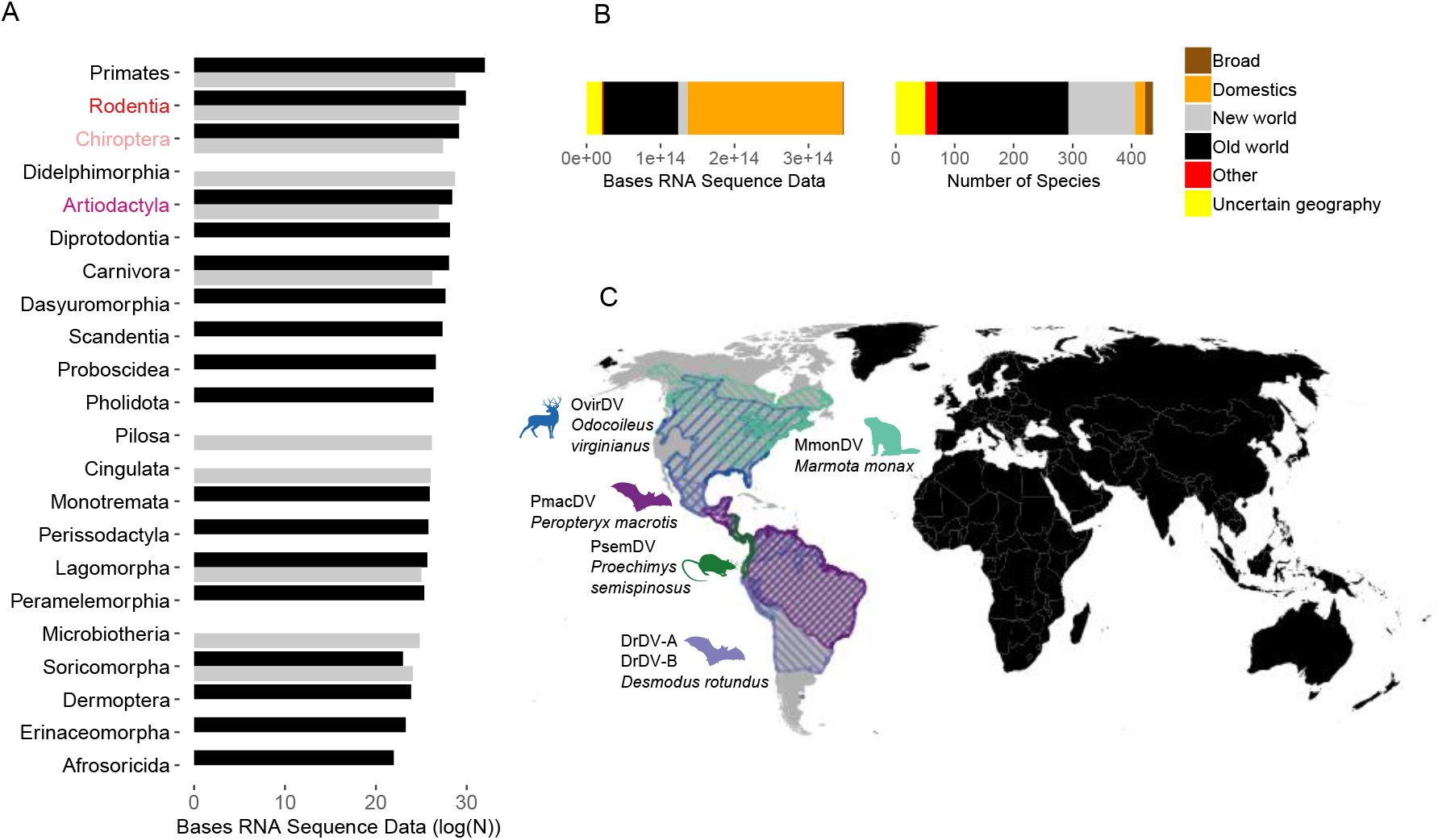
Geographic and taxonomic distribution of mammalian datasets and novel deltaviruses. (**A**) The host and geographic distribution of metagenomic and transcriptomic datasets searched for novel deltaviruses, note the log scale. Text colors indicate orders (red = Rodentia; pink = Chiroptera; Purple = Artiodactyla); bar colors indicate geography (black = Old World; gray = New World). (**B**) Stacked bar charts show the volume of mammalian datasets in units of RNA bases and the number of species searched, separated by species geography. Additional segments describe widely distributed domesticated animals (Domestics), datasets with genus-level metadata from broadly-distributed genera (Broad), datasets from cell lines or with taxonomic information only at the Class level (Other), and those which had no geographic range data available (Uncertain geography, mostly aquatic mammals). (**C**) Host distributions of newly discovered and recently reported deltaviruses, color coded by mammalian species (Data from IUCN). All except PsemDV were discovered through our search.

Searches revealed five deltaviruses spanning three mammalian orders: Artiodactyla (N=1), Chiroptera (N=3), and Rodentia (N=1; Fig. 1C). No deltaviruses were detected in non-human primates, indicating HDV as the sole known representative infecting the order Primates. Strikingly, despite over-representation of Old World-derived data by factors of (Artiodactyla), 5.8 (Chiroptera), 2.1 (Rodentia), all new mammalian deltaviruses originated from North and South American species (Fig. 1A,C and SI Appendix, Supplementary Results Section 1). Chiropteran deltaviruses included two genotypes from common vampire bats (*Desmodus rotundus*) which shared only 48.4-48.6% genome-wide nucleotide (nt) identity (hereafter, DrDV-A and DrDV-B; see SI Appendix, Fig. S1). A third deltavirus was identified in a liver transcriptome (accession SRR7910143; (12)) from a lesser dog-like bat (*Peropteryx macrotis*) from Mexico (PmacDV), but was more closely related to recently described deltaviruses from Tome’s spiny rat from Panama (PsemDV; (7)), sharing 95.9-97.4% amino acid (aa) and 93.0-95.7% nt identity. Additional genomes were recovered from transcriptomes derived from the pedicle tissue of white-tailed deer (*Odocoileus virginianus*; OvirDV; accession SRR4256033; (13)) and from a captive-born Eastern woodchuck (*Marmota monax*; MmonDV; accession SRR2136906; (14)). Bioinformatic screens recovered additional reads matching each genome in related datasets (either different individuals from the same study or different tissues from the same individuals), suggesting active infections (see SI Appendix, Table S2). All genomes had lengths 1669-1771 nt, high intramolecular base pairing, and contained genomic and antigenomic ribozymes characteristic of deltaviruses. The DrDV-A and DrDV-B genomes are more fully characterized in the SI Appendix (see SI Appendix, Fig. S2, Fig. S3, Table S3 and Supplementary Results Section 2). The other genomes and a case study on MmonDV infections in animals inoculated with woodchuck hepatitis virus are described by Edgar *et al*. (10).

Phylogenetic analysis of the small delta antigen (DAg) protein sequences using MrBayes (Fig. 2A) and a multi-species coalescent model in StarBeast (Fig. 2B) revealed multiple putative host shifts within the evolutionary history of mammalian deltaviruses. For instance, vampire bat deltaviruses were paraphyletic, suggesting at least two independent incursions into this species. Specifically, DrDV-A formed a clade with OvirDV and MmonDV (posterior probability, *PP* = 0.99/0.80 in MrBayes and StarBeast respectively) which was basal to HDV (*PP* = 0.65/0.81), while DrDV-B shared a most recent common ancestor with PmacDV and PsemDV (*PP* = 1/1). Rodent-associated deltaviruses (MmonDV and PsemDV) were also highly divergent and paraphyletic. Consequently, co-phylogenetic analyses using 1,000 randomly sampled topologies from StarBeast failed to reject independence of mammal and deltavirus phylogenies, consistent with a model of diversification by host shifting (Fig. 2B,C). Analyses of all deltavirus-host pairs (i.e. including highly divergent HDV-like agents) and an ‘ingroup’ clade containing mammalian along with avian and snake deltaviruses revealed somewhat greater dependence of the deltavirus phylogeny on the host phylogeny (see SI Appendix, Fig. S4). However, statistical significance varied across co-phylogenetic approaches and topological incongruences were evident among non-mammals, excluding co-speciation as the sole diversification process, even in deeper parts of the co-evolutionary history (see SI Appendix, Fig. S4, S5, Supplementary Results Section 3).

**Figure 2.**
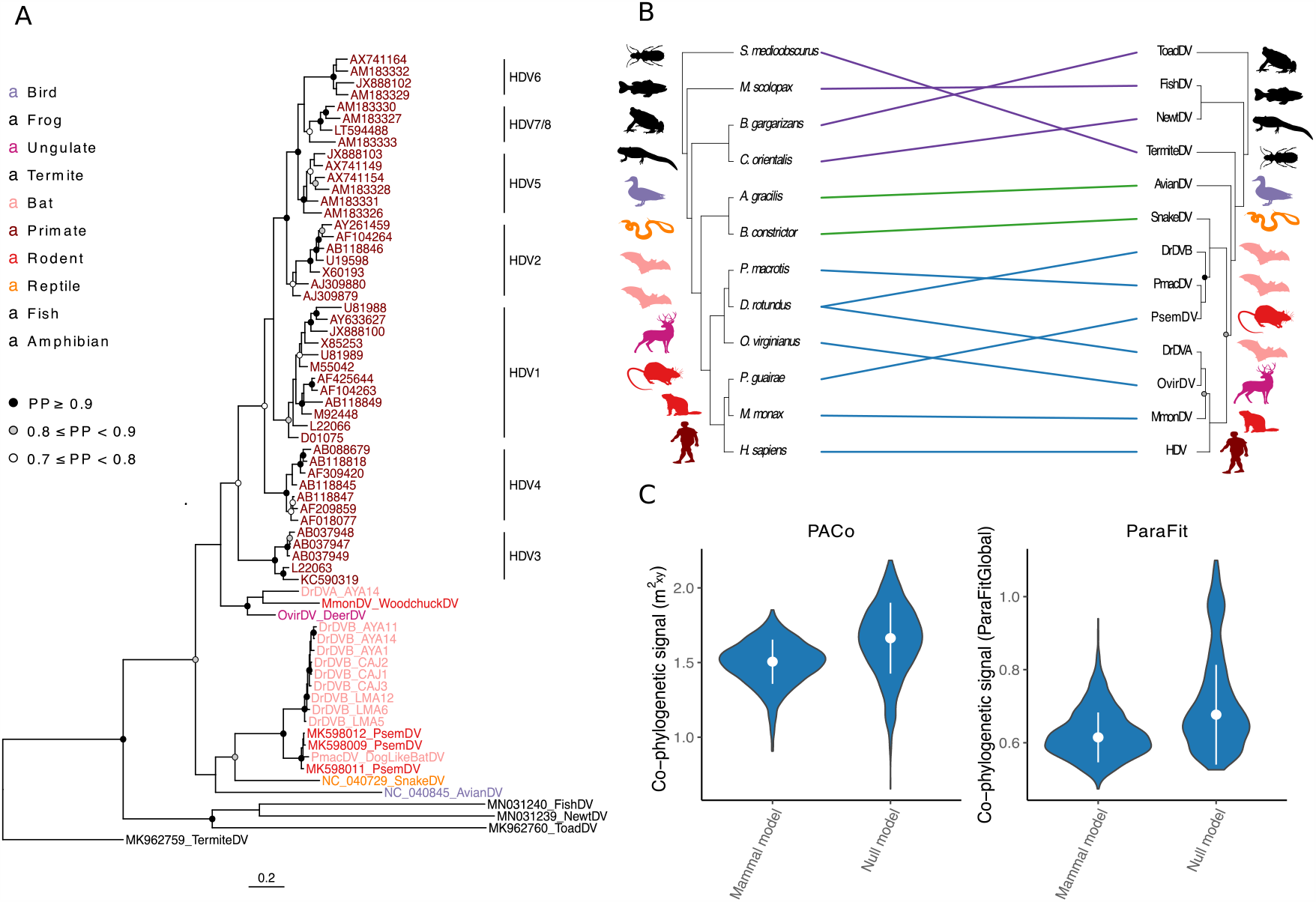
Evolutionary history of deltaviruses reveals host shifts among mammals. (**A**) Bayesian phylogeny of a 192 amino acid alignment of the DAg. Ingroup taxa including mammal, snake and avian deltaviruses are colored by order; other HDV-like taxa are shown in black. (**B**) Co-phylogeny depicting connections between the consensus deltavirus phylogeny from StarBeast and the host tree (from TimeTree.org). Links are colored according to subsets of data used in co-phylogenetic analyses; all taxa (purple + green + blue), ingroup (green + blue), or mammal (blue). Host taxa are: *Schedorhinotermes medioobscurus, Macroramphosus scolopax, Bufo gargarizans, Cynops orientalis, Anas gracilis, Boa constrictor, Peropteryx macrotis, Desmodus rotundus, Odocoileus virginianus, Proechimys guirae, Marmota monax*, and *Homo sapiens*. (**C**) Absence of phylogenetic dependence of the mammalian deltavirus phylogeny on the host phylogeny. Violin plots show distributions of test statistics from two co-phylogenetic approaches across 1,000 posterior trees relative to null models, along with medians and standard deviations. For PACo, higher values would indicate greater phylogenetic dependence; for ParaFit, lower values. Both approaches rejected a global model of co-speciation (*P* > 0.05).

Having extended the mammalian host range of deltaviruses to Neotropical bats, we subsequently explored the transmission dynamics, host range, and candidate helper virus associations within this group through a field study in three regions of Peru (Fig. 3A). Out of 240 *D. rotundus* saliva samples from 12 bat colonies collected in 2016-2017, RT-PCR targeting the DAg detected DrDV-A in single adult female from one of the two metagenomic pools that contained this genotype (bat ID 8299, site AYA14, see SI Appendix, Table S4, S5). In contrast, DrDV-B was detected in 17.1% of *D. rotundus* saliva samples (colony-level prevalence: 0-35%). Prevalence varied neither by region of Peru (Likelihood ratio test; χ^2^ = 3.21; d.f. = 2; *P* = 0.2) nor by bat age or sex (binomial generalized linear mixed model, Age: *P* = 0.38; Sex: *P* = 0.87), suggesting geographic and demographic ubiquity of infections. Given that vampire bats subsist on blood, deltavirus sequences encountered in bat saliva might represent contamination from infected prey. A small set of blood samples screened for DrDV-A (N = 60, including bat 8299) were negative. However, 6 out of 41 bats that were DrDV-B negative and 4 out of 18 bats with DrDV-B in saliva also contained DrDV-B in blood samples (Fig. 3B). In the 4 individuals with paired saliva and blood samples, DAg sequences were identical, supporting systemic infections. Significant spatial clustering of DrDV-B sequences at both regional and bat colony levels further supported local infection cycles driven by horizontal transmission among vampire bats (Fig. 3A and SI Appendix, Table S6).

**Figure 3.**
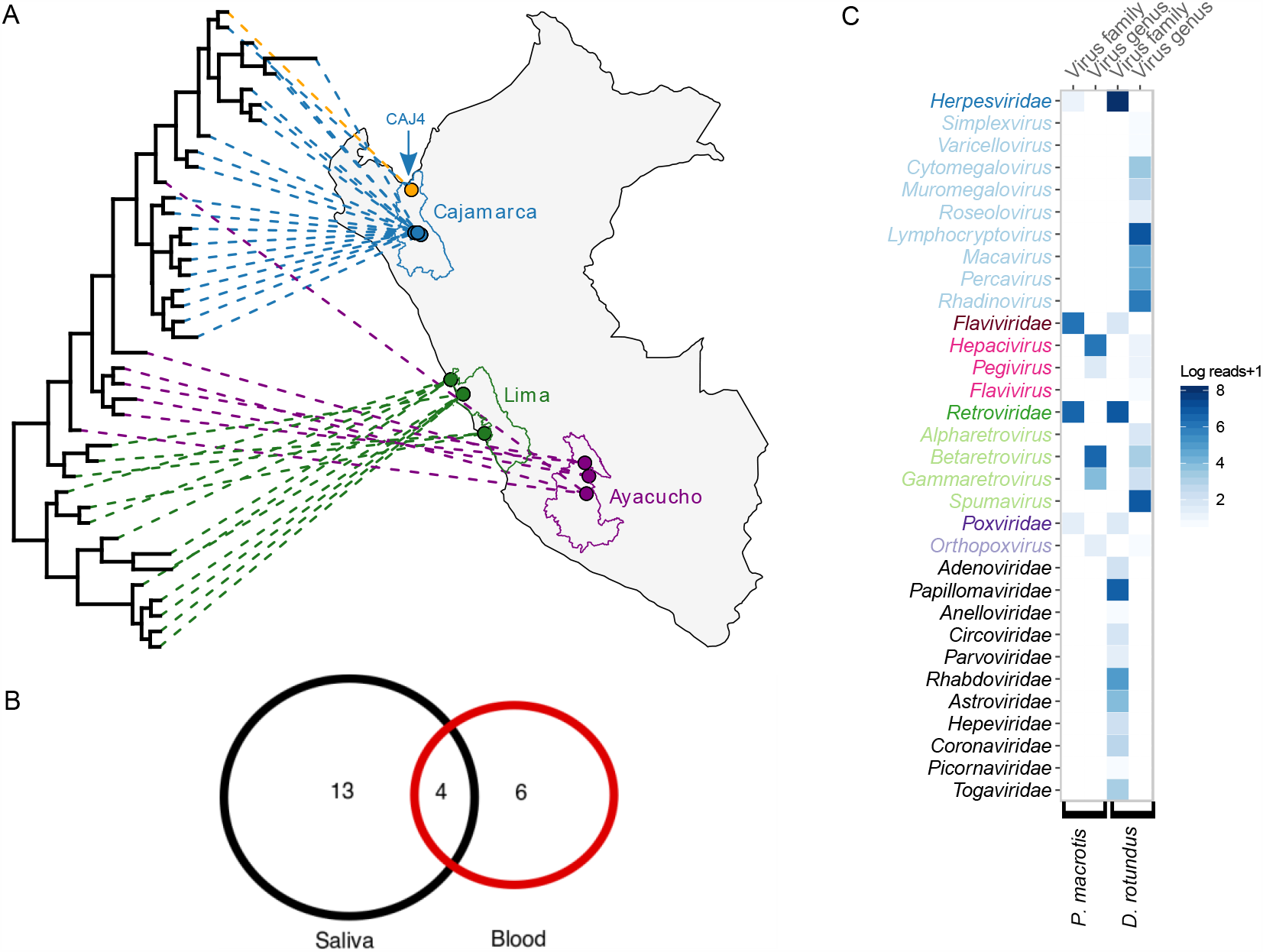
Transmission biology and candidate helper viruses for bat deltaviruses. (A) Bayesian phylogeny of a 214 nt alignment of DrDV-B DAg projected onto vampire bat capture locations in Peru. Lines and points are colored by administrative regions. Site CAJ4, where the *C. perspicillata* sequence was detected, is depicted in orange. (B) DrDV-B detections in saliva and blood. Numbers represent individual bats; the four bats in the center had genetically identical DrDV-B sequences in saliva and blood. (C) Mammal-infecting viral communities are shown for the *P. macrotis* liver transcriptome which contained PmacDV and combined *D. rotundus* saliva metagenomes. Viral taxa are colored by family, with lighter shades indicating genera within families, for overlapping viral families in both bat species. Viral families only present in one bat species are shown in black. Candidate helpers for OvirDV and MonDV are shown in SI Appendix, Fig. S9.

Given the evolutionary evidence for deltavirus host shifts, we hypothesized that spillover infections might also occur at detectable frequencies in sympatric Neotropical bats. Among 87 non-*D. rotundus* bats captured in or outside of *D. rotundus*-occupied roosts, RT-PCR detected deltavirus RNA in the saliva of a single Seba’s short-tailed bat (*Carollia perspicillata;* N=31 individuals; see SI Appendix, Fig. S6). This result was unlikely attributable to erroneous bat species assignment or laboratory contamination (see SI Appendix, Supplementary Results Section 4). The partial DAg recovered from the *C. perspicillata* was identical to a DrDV-B strain collected from a vampire bat in the same roost (CAJ4; Fig. 3A). Given the rapid evolution expected in deltaviruses (ca. 10^−3^ substitutions/site/year), genetic identity is most parsimoniously explained as horizontal transmission from *D. rotundus* to *C. perspicillata*, which was followed by an absence of (or short-lived) transmission among *C. perspicillata* at the time of sampling (15). This finding therefore demonstrates cross-species transmission on ecological timescales, a defining prerequisite for evolutionary diversification of deltaviruses through host shifting. Finally, we evaluated whether bat deltaviruses use hepadnavirus helpers akin to HDV (1). Consistent with a previous study, PCR screens of DrDV-positive and negative saliva (N=54) and blood samples (N=119), found no evidence of hepadnaviruses in vampire bats (16). To rule out divergent hepadnaviruses missed by PCR, we next used a bioinformatic pipeline to characterize viral communities in the metagenomic and transcriptomic datasets from deltavirus-infected bat species (Materials and Methods). Hepadnaviruses were again absent from all datasets (Fig. 3C). Together with the finding that all three bat-infecting deltavirus genomes lacked the farnesylation site thought to facilitate acquisition of the hepadnaviral envelope (see SI Appendix, Supplementary Results Section 2), use of hepadnavirus helpers by bat deltaviruses seems unlikely. To identify alternative plausible candidates, we quantified the abundance (approximated by sequence reads) of viral taxa that overlapped between the two deltavirus-infected bat species, *P. macrotus* and *D. rotundus*. In the *P. macrotus* liver which contained PMacDV, reads from hepaciviruses (*Flaviviridae*) spanned a complete viral genome (Genbank Third Party Annotation (TPA): BK013349) and outnumbered all other viral genera with the exception of Betaretroviruses, whose abundance may reflect endogenization in the host genome (Fig. 3C). Lower, but detectable, hepacivirus abundance in vampire bats may reflect the tissue tropism of these viruses or pooling of samples from multiple individuals. Intriguingly, hepaciviruses experimentally mobilize HDV *in vitro* and were found in 26 out of 30 PsemDV-infected rats (7, 17). Phylogenetic analysis of hepaciviruses associated with the deltavirus-positive host species (*P. macrotus, D. rotundus*, and *P. semispinosus*) revealed the candidate helpers to be highly divergent (see SI Appendix, Fig. S7), despite the apparently close relationships between the deltaviruses infecting these hosts. Reads matching *Poxviridae* formed small contigs in both libraries (*P. macrotus*: 229-1386 nt; *D. rotundus*: 358 nt) and particularly in *D. rotundus*, could not be excluded as false positives. Although non-opportunistic sampling is required to decisively identify the helpers of bat deltaviruses, existing evidence points to hepaciviruses as top contenders, perhaps using alternative enveloping mechanisms to HDV.

## Discussion

Unlike conventional pathogens (e.g., viruses, bacteria, protozoans), the obligatory dependence of deltaviruses on evolutionarily independent helpers creates a barrier to cross-species transmission that might be expected to promote host specificity. Data to test this hypothesis have been unavailable until now. Our study demonstrates transmission of deltaviruses among highly divergent mammalian orders on both ecological and evolutionary timescales.

Deltavirus host shifts could conceivably arise through several processes. Mobilization by non-viral micro-organisms (e.g., intra-cellular bacteria) is conceivable but has never been observed. Unaided spread through a yet undefined mechanism is also possible. However, given that the best-studied deltavirus (HDV) depends on viral envelopes to complete its life cycle and that conserved genomic features in related deltaviruses suggest a similar life history strategy, helper virus mediated host shifting is the most reasonable expectation. We and others have excluded hepadnavirus helpers for PmacDV, DrDVs, and PsemDV, yet natural HDV infections consistently involve HBV (1, 7). In light of this and the evidence presented here for host shifts among mammals, the contemporary HDV-HBV association must have arisen through acquisition of the hepadnaviral helper somewhere along the evolutionary divergence separating human and other mammal-infecting deltaviruses. Evidence that deltaviruses can exploit diverse enveloped viruses experimentally adds further weight to this conclusion (17, 18). As several new mammalian deltaviruses were detected with hepacivirus and poxvirus co-infections, either simultaneous host shifts of deltaviruses and helpers, or preferential deltavirus shifting among host species that are already infected with compatible helpers are plausible. If hepaciviruses are functioning as helpers, divergent phylogenetic relationships between species detected in bat and rodent hosts (see SI Appendix, Fig. S7) suggest they have been acquired as independently of deltaviruses rather than representing simultaneous host shifts or co-speciation of deltaviruses and helpers. Conclusively identifying the helper associations of novel mammalian deltaviruses and their evolutionary relationships will be crucial to disentangling these possibilities.

A limitation of our study was that the species in which novel deltaviruses were discovered were presumed to be definitive hosts (i.e. capable of sustained horizontal transmission). Consequently, some putative host shifts detected in our co-phylogenetic analysis may represent short-lived transmission chains in novel hosts or singleton infections, analogous to our observation of DrDV-B in *C. perspicillata*. For example, PmacDV from the Lesser dog-like bat clustered within the genetic diversity of PsemDV from rats, but infected two out of three individual bats analyzed, suggesting a recent cross-species transmission event followed by some currently unknown amount of onward transmission. Irrespective of the long-term outcomes of index infections, our results unequivocally support the conclusion that deltaviruses can transmit between divergent mammalian orders. The global distribution of deltavirus positive and negative datasets provides additional, independent evidence for host shifts. Even allowing for sub-detection due to variation in infection prevalence and dataset quality, deltaviruses should have been more widespread across the mammalian phylogeny than we observed (ca. 1% of species analyzed) if they co-speciated with their hosts. Given the presence of HDV in humans, non-human primates in particular would have been expected to host HDV-like deltaviruses in a co-speciation scenario, which was not observed. Moreover, the three deltavirus-infected mammalian orders occur in both the New and Old Worlds, yet non-human deltaviruses occurred exclusively in the Americas. Sampling biases cannot readily explain this pattern. By most metrics of sequencing effort, Old World mammals were over-represented each deltavirus-infected mammalian order, including at finer continental scales (see SI Appendix, Supplementary Results Section 1). Despite the large scale of our search, we evaluated < 10% of mammalian species. We therefore anticipate further discoveries of mammalian deltaviruses. Crucially, however, new viruses could not re-unite the paraphyletic rodent and bat deltaviruses or resolve widespread incongruence between mammal and deltavirus phylogenies, making our conclusions on host shifting robust.

The origin of HDV has been a longstanding mystery thwarted by the absence of closely related deltaviruses. The addition of 6 new mammalian deltaviruses by ourselves and others allowed us to re-evaluate this question (7, 10). The pervasiveness of host shifting among deltaviruses and our discovery of a clade of mammalian deltaviruses basal to HDV (albeit with variable support depending on phylogeny, *PP* = 0.64-0.81) strongly points to a zoonotic origin. Although the exact progenitor virus remains undiscovered, the exclusive detection of mammalian deltaviruses in New World species supports an “out of the Americas” explanation for origin and global spread of HDV (Fig. 1). The basal placement of the highly divergent Amazonian HDV genotype (HDV-3) within the phylogeny lends further credence to this scenario. It is therefore conceivable that mammal-infecting deltaviruses evolved in the Americas and the hypothesized arrival of HBV via human dispersal along the Beringian land bridge facilitated the zoonotic emergence of HDV (19). Though circumstantial, the earliest records of HDV are from the Amazon basin in the 1930s (20). The greater diversity of HDV genotypes outside of the Americas – long argued to support an Old World origin – may instead reflect diversification arising from geographic vicariance within human populations. We suggest that American mammals should be the focus of future efforts to discover the direct ancestor and zoonotic reservoir of HDV.

Our results show that deltaviruses jump between mammalian host species through an unusual process that most likely requires parasitizing evolutionarily independent viruses, and which has potential consequences for human and animal health. Since satellite viruses in general and HDV in particular tend to alter the pathogenesis and transmissibility of their helpers (21), our findings imply the potential for deltaviruses to act as host-switching virulence factors that could alter the progression of viral infections in multiple host species. The presence of deltaviruses in several mammalian orders, including in the saliva of sanguivorous bats which feed on humans, wildlife, and domestic animals, provides ecological opportunities for cross-species transmission to humans or other host species. Constraints on future host shifts are likely to differ from those of conventional animal pathogens. Specifically, given the broad cellular tropism of deltaviruses, interactions with viral helpers would likely be more important determinants of cross-species transmission than interactions with their hosts (17, 18). Consequently, anticipating future host shifts requires understanding the determinants and plasticity of deltavirus and helper compatibility along with the ecological factors that enable cross-species exposure.

## Materials and Methods

### 1. Virus discovery

#### a. Serratus

The Serratus platform was used to search published mammalian metagenomic and transcriptomic sequence datasets available in the NCBI Short Read Archive (SRA). Briefly, Serratus uses a cloud computing infrastructure to perform ultra-high throughput alignment of publicly available SRA short read datasets to viral genomes of interest (10). Due to the exceptional computational demands of this search and mutual interest between ourselves and another research team, Serratus searches were designed by both teams and carried out at the nucleotide and amino acid levels by Edgar *et al*. (10). Query sequences for Serratus searches included all HDV genotypes and all deltavirus-like genomes which were publicly available at the time of the search (July 2020) along with representative genomes from DrDV-A and DrDV-B, which our team had already discovered. Results were shared among the two teams who subsequently pursued complementary lines of investigation. Novel mammalian deltaviruses were discovered using nucleotide (OvirDV) and amino acid level (Mmon DV and PmacDV) searches of the mammalian SRA search space.

#### b. Neotropical bat metagenomic sequencing

Total nucleic acid was extracted from archived saliva swabs from Neotropical bats on a Kingfisher Flex 96 automated extraction machine (ThermoFisher Scientific) with the Biosprint One-for-all Vet Kit (Qiagen) using a modified version of the manufacturer’s protocol as described previously (11). Ten pools of nucleic acids from vampire bats and other bat species were created for shotgun metagenomic sequencing (see SI Appendix, Table S1). Eight pools comprised samples from bats in the same genus (2-10 individuals per pool depending on availability of samples, 30 μL total nucleic acid per individual). The CAJ1_SV vampire bat pool from (22) which contained deltavirus reads was included as a sequencing control. The final pool (“Rare species”) comprised 8 other bat species that had only one individual sampled each. Pools were treated with DNAse I (Ambion) and purified using RNAClean XP beads (Agencourt) following (11). Libraries were prepared using the SMARTer Stranded Total RNA-Seq Kit v2 - Pico Input Mammalian (Clontech) and sequenced on an Illumina NextSeq500 at The University of Glasgow Polyomics Facility. Samples were bioinformatically processed for viral discovery as described previously (11), with a slight modification to the read trimming step to account for shorter reads and a different library preparation kit.

#### c. DrDV genome assembly and annotation

DrDV genomes were assembled using SPAdes (23) and refined by mapping cleaned reads back to SPAdes-generated contigs within Geneious v 7.1.7 (24). Regions of overlapping sequence at the ends of genomes due to linear *de novo* assemblies of circular genomes were resolved manually. Genome circularity was confirmed based on the presence of overlapping reads across the entire circular genome of both DrDVs. The amino acid sequence of the small delta antigen protein (DAg) was extracted from sequences using getorf (25). Other smaller identified open reading frames did not exhibit significant homology when evaluated by protein blast against Genbank. Nucleotide sequences of full deltavirus genomes and amino acid sequences of DAg were aligned along with representative sequences from other deltaviruses using the E-INS-i algorithm in MAFFT v 7.017 (26). Genetic distances as percent identities were calculated based on an untrimmed full genome alignment of 2,321nt and an untrimmed delta antigen alignment of 281aa. Protein domain homology of the DAg was analyzed using Hhpred (27). Ribozymes were identified manually by examining the region upstream of the delta antigen open reading frame where ribozymes are located in other deltavirus genomes (4, 5). RNA secondary structure and self-complementarity were determined using the webservers for mFold (28) and RNAStructure (29). We found no evidence of recombination in nucleotide alignments of DrDV DAg according to the program GARD (30) on the Datamonkey webserver (31). Genome assembly and annotation of PmacDV, OvirDV, and MmonDV are described in (10). We used blast searches of novel mammalian deltavirus genomes (blastn) and deltavirus antigen protein sequences (tblastn) against published host genomes on Genbank to evaluate the possible presence of endogenous deltavirus-like elements. DrDV-A and DrDV-B sequences were searched against the *Desmodus rotundus* genome assembly (GCA_002940915.2), MmonDV sequences were searched against the *Marmota monax* genome assembly (GCA_901343595.1), and OvirDV sequences were searched against the *Odocoileus virginianus* RefSeq genome (NC_015247.1). There was no published genome of *Peropteryx macrotis* available on Genbank so we used blastn and tblastn to compare the genome and delta antigen protein sequence against all of Genbank, restricting results by organism *P. macrotis* (taxid:249015). None of these searches yielded any hits, suggesting there is currently no evidence these deltaviruses have endogenized in their hosts.

#### d. Evaluation of deltavirus positive cohorts

To establish that deltaviruses were likely to be actively infecting hosts in which they were detected, and not laboratory contamination or incidental detection of environmentally derived RNA, we searched for evidence of deltavirus infections in additional samples from the various studies that detected full genomes. Samples included sequencing libraries derived from both different individuals in the same study and different tissues from the same individuals. Searches used bwa (32) to map raw reads from deltavirus positive cohorts to the corresponding novel deltavirus genomes which had been detected in those same cohorts. Genome remapping was performed for all vampire bat libraries, two other *Peropteryx macrotis* libraries and all other Neotropical bat species sequenced in the same study (12) and other pooled tissue samples sequenced by RNASeq from *Odocoileus virginianus* in the same study (13). Results are shown in SI Appendix, Table S2. Deltavirus reads from additional individuals and timepoints from the *Marmota monax* study (14) are described in (10).

#### e. Global biogeographic analysis of deltavirus presence and absence

To characterize the global distribution of mammal-infecting deltaviruses, we used the metadata of each SRA accession queried in the Serratus search to identify the associated host. We focused primarily on the mammalian dataset, which was generated by the SRA search query (“Mammalia”[Organism] NOT “Homo sapiens”[Organism] NOT “Mus musculus”[Organism]) AND (“type_rnaseq”[Filter] OR “metagenomic”[Filter] OR “metatranscriptomic”[Filter]) AND “platform illumina”[Properties]. All analyses were performed in R version 3.5.1 (33). We removed libraries with persistent errors which had not completed in the Serratus search. For remaining libraries, when host identification information was available to the species level, we matched Latin binomial species names to Pantheria, a dataset which contains the centroids of mammalian geographic distributions, and used an R script to assign species to continents using these geographic data (34). Subspecies present in scientific names of SRA meta-data were re-assigned to species level and recently updated binomial names were changed to match the Pantheria dataset. Due to the fact that mammalian taxonomic data in Pantheria date to 2005, some former Orders which are no longer in use (e.g. Soricomorpha) appear in our data but are not expected to affect the results of analyses. Forty-eight species whose centroids occurred in water bodies were assigned to continents by manually inspecting species ranges. Widely-distributed domesticated animals, datasets with genus-level metadata from broadly-distributed genera, datasets from cell lines or with taxonomic information only at the Class level, and those which had no geographic range data available (mostly aquatic mammals) were searched by Serratus, but excluded from geographic comparisons. We quantified geographic and taxonomic biases in our dataset both in units of bases of RNA sequenced and number of species investigated.

Although most mammalian metagenomic and transcriptomic libraries were captured in the mammalian search, we also examined datasets from SRA search queries for vertebrates, metagenomes, and viromes to ensure that all relevant libraries were captured in our measures of search effort. For these datasets, we removed libraries with persistent errors and calculated search effort as bases of RNA sequenced across the three orders in which deltaviruses were discovered, removing libraries named for specific viral taxa which may have been enriched for these taxa and therefore do not represent a likely source of novel viruses. As these libraries lacked species-level meta-data (hence their exclusion from the mammalian search above), we could not systematically calculate number of species in these datasets. Additional search queries are available at https://github.com/ababaian/serratus/wiki/SRA-queries.

### 2. Phylogenetic analyses

#### a. Bayesian phylogeny using MrBayes

Phylogenetic analysis was performed on complete DAg amino acid sequences to place mammalian deltaviruses relative to HDV and other described deltaviruses. Representative sequences from each clade of HDV and other previously published deltaviruses were aligned with DrDVs (sequences generated by RT-PCR, described in Methods section 3b), PmacDV, OvirDV and MmonDV using the E-INS-i algorithm and JTT2000 scoring matrix of MAFFT within Geneious. Large delta antigen sequences from HDV were trimmed manually to small delta antigen length, and the alignment was further trimmed in trimal using the automated1 setting to a final length of 192aa. The best substitution model (JTTDCMut+F+G4) was determined using ModelFinder (35) within IQTree 2 (36). Bayesian phylogenetic analysis was performed using the most similar model available within MrBayes (JTT+F+G4). The analysis was run for 5,000,000 generations and sampled every 2,500 generations, with the first 500 trees discarded as burn-in to generate the consensus tree.

#### b. Bayesian phylogeny using StarBeast

We used StarBeast to generate a species-level phylogeny for the co-phylogenetic analysis, using the same amino acid alignment of complete DAg which was used in the MrBayes analysis. StarBeast is typically used with multi-locus sequence data from multiple individuals per species but can also be applied to single gene alignments (37). Notably, a preliminary analysis suggested this approach was more conservative than a constant effective size coalescent model in BEAST which substantially inflated posterior probabilities on nodes across the tree relative to the MrBayes analysis (Fig. 2A). The StarBeast multi-species coalescent analysis was carried out as two duplicate runs of 50 million generations (sampling every 5000 generations) in BEAST2, using the JTT+G substitution model, the linear with constant root model for the species tree population size, and a Yule speciation model. Combined log files were assessed for convergence and effective sample size values >200 using TRACER. Twenty percent of trees were discarded as burn-in prior to generating the consensus tree.

#### c. Co-phylogeny

Co-phylogenetic analyses were performed in R using PACo (38, 39) and ParaFit (40, 41). Analyses were performed on three subsets of matched host-deltavirus data: all taxa, ingroup taxa (mammals, bird and snake) and mammals only. Host datasets consisted of distance matrices derived from a TimeTree phylogeny (timetree.org). For metagenomic libraries which contained individuals of multiple species (AvianDV and FishDV), one host was selected for inclusion (*Anas gracilis* and *Macroramphosus scolopax*, respectively). Host data were not available in TimeTree for two species in which deltaviruses were discovered (*Proechimys semispinosus* and *Schedorhinotermes intermedius*), so available congeners were substituted (*Proechimys guairae* and *Schedorhinotermes medioobscurus*, respectively). Virus datasets consisted of distances matrices from posterior species trees generated in StarBeast. Co-phylogeny analyses performed using virus distance matrices derived from posterior MrBayes trees, pruned to contain only relevant taxa, yielded qualitatively similar results. For both analyses, the principal coordinates analysis of distance matrices was performed with the ‘cailliez’ correction. Since units of branch length differed between host and virus trees, all distance matrices were normalized prior to co-evolutionary analyses. To account for phylogenetic uncertainty in the evolutionary history of deltaviruses, analyses were carried out using 1,000 trees randomly selected from the posterior distribution of the Bayesian phylogenetic analyses (separately for StarBeast and MrBayes). Due to uncertain placement of HDV3 in both phylogenies, one representative of HDV was randomly selected for each iteration. For each tree, we calculated summary statistics (see below) describing the dependence of the deltavirus phylogeny on the host phylogeny. *P*-values were estimated using 1,000 permutations of host-virus associations.

For PACo analyses, the null model selected was r0, which assumes that virus phylogeny tracks the host phylogeny. Levels of co-phylogenetic signal were evaluated as the median global sum of squared residuals (m^2^_xy_) and mean significance (*P*-values), averaged over the 1,000 posterior trees. Empirical distributions were compared to null distributions for each dataset. For ParaFit analyses, the levels of co-phylogenetic signal in each dataset were evaluated as the median sum of squares of the fourth corner matrix (ParaFitGlobal) and mean significance (*P*-values), averaged over 1,000 posterior trees. ParaFit calculates the significance of the global host-virus association statistic by randomly permuting hosts in the host-virus association matrix to create a null distribution. Since Parafit does not provide this distribution to users, we approximated it for visualization by manually re-estimating the global host-virus association statistic for 1,000 random permutations of hosts in the host-virus association matrix. Phylogenies and co-phylogenies were visualized in R using the packages ‘ape’ (41), ‘phangorn’ (42), ‘phytools’ (43), and ‘ggtree’ (44).

### 3. Deltaviruses in Neotropical bats

#### a. Capture and sampling of wild bats

To examine DrDV prevalence in vampire bats, we studied 12 sites in three departments of Peru between 2016-2017 (Fig. 3A). Age and sex of bats were determined as described previously (11). Saliva samples were collected by allowing bats to chew on sterile cotton-tipped wooden swabs (Fisherbrand). Blood was collected from vampire bats only by lancing the propatagial vein and saturating a sterile cotton-tipped wooden swab with blood. Swabs were stored in 1 mL RNALater (Ambion) overnight at 4°C before being transferred to dry ice and stored in −70°C freezers.

Bat sampling protocols were approved by the Research Ethics Committee of the University of Glasgow School of Medical, Veterinary and Life Sciences (Ref081/15), the University of Georgia Animal Care and Use Committee (A2014 04-016-Y3-A5), and the Peruvian Government (RD-009-2015-SERFOR-DGGSPFFS, RD-264-2015-SERFOR-DGGSPFFS, RD-142-2015-SERFOR-DGGSPFFS, RD-054-2016-SERFOR-DGGSPFFS).

#### b. RT-PCR and sequencing of blood and saliva samples

Primers were designed to screen bat saliva and blood samples for a conserved region of the DAg protein of DrDV-A (236nt) and DrDV-B (231nt), by hemi-nested and nested RT-PCR respectively (see SI Appendix, Table S4). Alternative primers were designed to amplify the complete DAg for DrDV-A (707nt) and DrDV-B (948nt) using a one-step RT-PCR (see SI Appendix, Table S4). We used RT-PCR to screen vampire bat saliva samples (described in Methods section 3a), as well as saliva samples from additional Neotropical bat species included in metagenomic sequencing pools (described in Methods section 1b) and further archived samples from non-vampire bat species which were withheld from metagenomic pools in order to balance pool sizes (see SI Appendix, Fig. S6). A subset of vampire bat blood samples were also screened by RT-PCR; blood samples were unavailable from non-vampire bat species. cDNA was generated from total nucleic acid extracts using the Protoscript II First Strand cDNA synthesis kit with random hexamers; RNA and random hexamers were heated for 5 minutes at 65°C then placed on ice. Protoscript II reaction mix and Protoscript II enzyme mix were added to a final concentration of 1x, and the reaction was incubated at 25°C for 5 minutes, 42°C for 15 minutes, and 80°C for 5 minutes. PCR was performed using Q5 High-Fidelity DNA Polymerase (NEB). Each reaction contained 1x Q5 reaction buffer, 200 μM dNTPs, 0.5 μM each primer, 0.02 U/μL Q5 High Fidelity DNA polymerase and either 2.5 μL cDNA or 1 μL Round 1 PCR product. Reactions were incubated at 98°C for 30 seconds, followed by 40 cycles of 98°C for 10 seconds, 61-65°C for 30 seconds (or 58-60°C for 30 seconds for the complete DAg), 72°C for 40 seconds, and a final elongation step of 72°C for 2 minutes. PCR products of the correct size were confirmed by re-amplification from cDNA or total nucleic acid extracts and/or Sanger sequencing (Eurofins Genomics).

#### c. Bat species confirmation

We confirmed the morphological species assignment of the *C. perspicillata* individual in which DrDV-B was detected by sequencing cytochrome B. Cytochrome B was amplified from the same saliva sample in which DrDV-B was detected using primers Bat 05A and Bat 04A (45) and GoTaq Green Master Mix (Promega) according to the manufacturer’s instructions, and the resulting product was Sanger sequenced (Eurofins Genomics) then evaluated by nucleotide blast against Genbank.

#### d. Genetic diversity and distribution of DrDV-B

To examine relationships among DrDV-B sequences, Bayesian phylogenetic analysis was performed on a 214nt fragment of the DAg. Sequences from saliva and blood of 41 *D. rotundus* and saliva from one *C. perspicillata* were aligned using MAFFT within Geneious. Duplicate sequences originating from the blood and saliva of the same individuals were removed. Alignments were trimmed using Trimal (46) with automated1 settings, and the best model of sequence evolution was determined using jModelTest2 (47). Phylogenetic analysis was performed using MrBayes 3.6.2 (48) with the GTR+I model. The analysis was run for 4,000,000 generations and sampled every 2,000 generations, with the first 1,000 trees removed as burn-in. The association between phylogenetic relationships and location at both the regional and colony level was tested using BaTS (49) with 1,000 posterior trees and 1,000 replicates to generate the null distribution.

#### e. Statistical analyses of DrDV-B

We modeled the effects of age and sex on DrDV-B presence in saliva using a binomial generalized linear mixed model (GLMM) in the package lme4 (50) in R. Age (female/male) and sex (adult/subadult) were modeled as categorical variables, with site included as a random effect. We also evaluated differences in DrDV-B prevalence between regions of Peru using a binomial generalized linear model (GLM), and used the *Anova* function of the *car* package (51) to calculate the likelihood ratio χ^2^ test statistic.

### 4. Identifying candidate helper viruses for mammalian deltaviruses

#### a. Hepadnavirus screening in vampire bats

We tested samples for the presence of bat hepadnavirus as a candidate helper virus to DrDV. DNA from saliva and blood samples was screened for HBV-like viruses using pan-*Hepadnaviridae* primers (HBV-F248, HBV-R397, HBV-R450a, HBV-R450b; see SI Appendix, Table S4) and PCR protocols (16). We used a plasmid carrying a 1.3-mer genome of human HBV that is particle assembly defective but replication competent as a positive control.

#### b. Bioinformatic screening of published metagenomic datasets

We performed comprehensive virus discovery using an in-house bioinformatic pipeline (11) on sequence datasets containing deltaviruses to identify candidate helper viruses. Datasets analyzed included all vampire bat datasets (22 from (11), 46 from (22)), *P. macrotis* datasets (SRR7910142, SRR7910143, SRR7910144), *O. virginianus* datasets (SRR4256025-SRR4256034), and *M. monax* datasets (SRR2136906, SRR2136907). Briefly, after quality trimming and filtering, reads were analyzed by blastx using DIAMOND (52) against a RefSeq database to remove bacterial and eukaryotic reads. Remaining reads were then *de novo* assembled using SPAdes (23) and resulting contigs were analyzed by blastx using DIAMOND against a non-redundant (NR) protein database (53). KronaTools (54) and MEGAN (55) were used to visualize and report taxonomic assignments.

## Acknowledgments

We thank Jaime Pacheco, Luigi Carrasco, Yosselym Luzon, Saori Grillo and Megan Griffiths for field and laboratory assistance; Megan Griffiths, Joseph Hughes, and Matt Hutchinson for analysis advice; and Ana da Silva Filipe, Felix Drexler, Pablo Murcia and Mafalda Viana for comments on earlier versions of the manuscript. We thank the Serratus team, particularly Artem Babaian and Robert Edgar for assistance with Serratus.

## Data and materials availability

DrDV genome sequences are available on Genbank (accessions MT649206-MT649209). PmacDV, OvirDV and MmonDV genome sequences are available at https://serratus.io/access. The *Peropteryx macrotis* Hepacivirus genome sequence is available in the Third Party Annotation Section of the DDBJ/ENA/GenBank databases under the accession number TPA: BK013349. Vampire bat hepacivirus contigs are available on Genbank (accession numbers MW249008 and MW249009). Peruvian bat metagenomes are available in ENA project PRJEB35111. Scripts used for bioinformatic analyses are available on GitHub (https://github.com/rjorton/Allmond).

## Supplementary Information

### Supplementary Results

#### 1. Evaluation of continent level geographic biases in Serratus data

We sought to confirm that the sole detection of mammalian deltaviruses in the Americas in three different mammalian orders was unlikely to have arisen from sampling biases in the RNA sequence datasets that we analyzed. The main text illustrates geographic patterns at broad scales and shows that New World species were under-represented relative to Old World species, indicating that sampling biases were unlikely to explain the absence of deltaviruses from Old World species. However, we also examined biases at finer geographic and taxonomic scales, focusing on the mammalian SRA search query dataset for which there was species and continent level information. At the continent level, data volumes (in bases of RNA sequenced) declined from Asia (6.35e13), Africa (2.89e13), South America (7.99e12), North America (4.93e12), Europe (4.86e12), to Australia (3.75e12) (Fig S8). This implies that Africa (the previously assumed origin of HDV) has more than double the data of the Americas combined. Similar patterns were evident within the deltavirus-infected mammalian orders. For Artiodactyla, North and South American datasets were ranked 3^rd^ and 5^th^ respectively among the 5 continents which had sequence data. Although there was less Artiodactyla data from Africa than North America, Asia (1^st^) and Europe (2^nd^) had 2 and 1.6-fold more data than the Americas combined. For bats, North and South American datasets were ranked 5^th^ and 6^th^ among the 6 continents with data, and Europe (1^st^) and Africa (2^nd^) had 2.5 and 1.2-fold more data than datasets combined across the Americas. For rodents, North America datasets were ranked 2^nd^, while South American datasets were ranked 5^th^ out of the 5 continents with data, but Asia (1^st^) had 1.1-fold more data than the Americas combined and Africa had 3.6 times more data than South America.

We also examined potential biases in search effort by number of species sequenced per continent. There were equal or fewer species of Artiodactyla sequenced in North (N = 5) and South America (N=1) compared to Europe (N = 5), Asia (N = 8) and Africa (N = 14). Similarly, rodent species sampled from Old World continents (N = 48; Europe [N = 8], Africa [N = 14], and Asia [N = 26]) outnumbered those in the New World (N = 33; North America [N = 21], South America [N = 12]. Neither Rodentia nor Artiodactyla datasets were searched from Australia. In contrast, there were more bat species in North and South American datasets (N = 4 and 42, respectively) compared to Asia (N = 22), Africa (N = 5), Europe (N = 2), and Australia (N =1) leading to a slight bias toward New World bats (46 species vs 30 Old World species). Consequently, in the ‘mammalian’ dataset, New World bats were more numerous at the species level, but had fewer individuals tested per species and/or less sequencing depth per species.

We further examined datasets generated by related search queries (vertebrate, metagenome and virome) which included some libraries from mammals that were excluded from the ‘mammalian’ dataset (see Materials and Methods, Section 1e). The vertebrate dataset contained no matches to the three mammalian Orders of interest (Rodentia, Artiodactyla, Chiroptera). For Rodentia, the metagenome dataset contained 380 libraries (9.7e7 bases) identified as “mouse metagenome” and the virome dataset contained 171 libraries (1e11 bases) which were identified as “mouse gut metagenome” or “rodent metagenome” or “Rattus” or “rat gut metagenome”. These could not be assigned geographic provenance and likely represented laboratory animals. For Artiodactyla, the virome dataset contained 317 libraries (1e11 bases) which were identified as “pig gut metagenome” or “pig metagenome” or “bovine gut metagenome”. As these libraries likely derived from either experimental or domestic animals, we conclude that additional data from these two Orders are unlikely to influence geographic or taxonomic bias. For Chiroptera, there were a total of 25 libraries (2.4e10 bases) identified as “bat metagenome”. The vast majority (N=24) were from Old World bats. Eleven derived from a study of bat rotaviruses (PRJNA562472) which of which 10 were collected from Old world locations (Ghana, Bulgaria) and one from New World (Costa Rica). Two libraries were bat viral metagenomes generated from samples collected in South Africa (SRR5889194; SRR5889129), and twelve libraries were generated from bats sampled in China (PRJNA379515). There were 14 bat species analyzed in all of these libraries, of which eight were not included in the mammalian SRA dataset, bringing the total number of Old World bat species to 38. Therefore, by the metric of number of species, New World bats remained slightly over-represented (38 versus 46 species) though as mentioned above, Old World bat derived datasets were sequenced more comprehensively and covered a larger number of continents.

Overall, across the three orders in which we detected deltaviruses, fewer species were studied in North and South American datasets (85 species) compared to those from Africa, Asia, Europe and Australia (105 species) and the total volume of RNA sequenced was 2.7 times greater for Old world species (1.64e13 bases RNA) than New world species (6.02e12). We therefore conclude that the exclusive presence of deltaviruses in American mammals is unlikely to represent geographic biases in our datasets.

#### 2. Large delta antigen in novel mammalian deltaviruses

In HDV, the large delta antigen protein (L-HDAg) is produced by RNA editing of the UAG stop codon to include 19 additional aa (1) and contains a farnesylation site which interacts with HBV (2). The DrDV-B DAg from the genome from bat colony CAJ1 terminated in UAG, which if edited similarly to HDV would generate a putative L-DAg containing an additional 28 aa (Fig. S3). In contrast, DrDV-B DAg from the two other bat colonies from which genomes were sequenced (LMA6 and AYA11), as well as DrDV-A DAg, terminated in a UAA stop codon so would not appear to be similarly edited, although it is possible to extend the open reading frames through frameshifting (3). Importantly, no putative vampire bat L-DAg generated through either RNA editing or frameshifting contained a farnesylation site. PmacDV, OvirDV, and MmonDV also did not contain apparent L-DAg extensions or farnesylation sites (4).

#### 3. Co-phylogenetic results across posterior distributions of trees from two Bayesian searches and two different co-phylogeny analyses

Consensus topologies differed slightly between the phylogenetic analyses of deltaviruses using a multi-species coalescent model in StarBeast and a coalescent model in MrBayes, particularly in relation to the termite-associated deltavirus-like agent and avian deltavirus (compare Fig. 2A and 2B). We therefore repeated our co-phylogenetic analysis using 1,000 trees from the MrBayes analysis to verify that our conclusions were robust to this topological inconsistency. Analyses performed using virus distance matrices derived from posterior MrBayes trees were congruent with those in StarBeast.

In the broadest analyses of all taxa, results differed slightly among co-phylogenetic analyses. Specifically, PACo analyses supported the dependence of the deltavirus phylogeny on the host phylogeny (StarBeast trees: m^2^_xy_ = 0.57 (standard deviation = 0.48-0.66); m^2^_xy_null_ = 1.27 (1.11-1.44); *P* = 0.01; MrBayes trees: (m^2^_xy_ = 0.53 (0.46-0.6); m^2^_xy_null_ = 1.26 (1.1-1.42); *P* = 0.002). However, ParaFit analyses supported independence of the virus and host phylogenies based on both StarBeast trees (ParaFitGlobal = 0.72 (standard deviation = 0.59-0.86); *P*= 0.09) and MrBayes trees (ParaFitGlobal = 0.63 (0.52-0.75); *P*= 0.07). Similarly, for ingroup taxa (mammalian, avian and snake deltaviruses), PACo detected evidence of co-phylogeny using both sets of trees (Starbeast: m^2^_xy_ = 0.81 (0.66-0.96); m^2^_xy_null_ = 1.3 (1.14-1.46); *P* = 0.02; MrBayes: m^2^_xy_ = 0.76 (0.69-0.83); m^2^_xy_null_ = 1.28 (1.1-1.46); *P* = 0.01), while ParaFit analyses with StarBeast trees (ParaFitGlobal = 0.8 (0.7-0.89); *P*= 0.12) and MrBayes trees (ParaFitGlobal = 0.68 (0.62-0.75); *P*= 0.09) found no significant support for co-phylogeny.

All analyses of mammalian deltaviruses failed to reject the null hypothesis of independence of phylogenies. These results were consistent when with both StarBeast and MrBayes trees in PACo (StarBeast: m^2^_xy_ = 1.51 (1.36-1.65); m^2^_xy_null_ = 1.66 (1.43-1.9); *P* = 0.28; MrBayes: m^2^_xy_ = 1.16 (1.07-1.25); m^2^_xy_null_ = 1.22 (0.97-1.48); *P* = 0.35), as well as in ParaFit (StarBeast: ParaFitGlobal = 0.61 (0.55-0.68); *P*= 0.52; MrBayes: ParaFitGlobal = 0.49 (0.44-0.54); *P*= 0.5).

In summary, both PACo and ParaFit analyses of StarBeast and MrBayes trees showed no support for co-phylogeny in the mammalian dataset. Including more divergent host-virus pairs increased support for co-phylogeny in the all taxa and ingroup datasets, with these results being statistically significant in PACo analyses but not in ParaFit analyses. Inconsistent support for phylogenetic independence at broader scales may reflect variation in the sensitivity of different analyses to detect phylogenetic congruence which occurs in only a subset of branches. For example, the non-ingroup deltavirus-like agents formed a polytomy of long branches and were found in the most divergent hosts from mammals, which may have inflated co-phylogenetic signal (Fig. S5). Regardless, given the consistent evidence against a co-speciation model among mammals and incongruences observed among other taxa in the consensus topologies, these findings illustrate that a model of co-speciation alone cannot explain the evolutionary relationships of deltaviruses and their hosts.

#### 4. Putative cross-species transmission of DrDV-B to a frugivorous bat

The detection of a vampire bat associated deltavirus in a frugivorous bat (*Carollia perspicillata)* is strongly suggestive of cross-species transmission but might also arise through mis-assignment of bat species in the field or contamination of samples during laboratory processing. To exclude the possibility of host species mis-identification, we confirmed morphological species assignment by sequencing Cytochrome B from the same saliva sample in which we amplified deltavirus (see Methods), which showed 99.49% identity with a published *C. perspicillata* sequence in Genbank (Accession AF511977.1). Laboratory contamination was minimized by processing all samples through a dedicated PCR pipeline with a one directional workflow. PCR reagents are stored and master mixes prepared in a laboratory that is DNA/RNA free, and which cannot be entered after going into any other lab. Field collected samples from bats are extracted and handled in a room strictly used for clinical samples which cannot be entered after going in any other lab aside from the master mix room. To further exclude laboratory contamination, we independently amplified the *C. perspicillata* deltavirus product from two separate batches of cDNA. We used only round 1 primers of a nested PCR to avoid detecting trace amounts of potential contamination; in vampire bats only 68% of individuals deemed positive after round 2 were also positive in round 1. Furthermore, in the laboratory, samples from other bat species were handled separately from samples collected from vampire bats, with extractions and PCRs being performed on different days. As discussed in the main text, the absence of genetic divergence from sympatric strains in *D. rotundus* indicates limited or no onward transmission of DrDV-B in *C. perspicillata*. Whether the *C. perspicillata* sustained an actively replicating infection is uncertain, although detection in a single round of PCR (which was true for only 68% of DrDV-positive vampire bats) implies an intensity of infection which could suggest DrDV replication in the recipient host, though this would require further testing to confirm. Definitively resolving the extent of DrDV-B replication could be achieved using a quantitative RT-PCR targeting the DrDV antigenome. Such assays do not currently exist and after the confirmatory testing above, in addition to metagenomic sequencing, we unfortunately would no longer have sufficient RNA available from the *C. perspicillata* bat to run such a test if it were available. In summary, we are confident that the individual in which the deltavirus was detected is a *C. perspicillata* and we believe the most likely explanation to be cross-species transmission in nature, though whether this represents an active infection remains uncertain.

#### 5. Candidate helper viruses of OvirDV and MmonDV

We also examined viral communities in *O. virginianus* and *M. monax* libraries for candidate helpers. Given that MmonDV was detected in animals experimentally inoculated with *Woodchuck hepatitis virus* (WHV, a hepadnavirus), these libraries were unsurprisingly dominated by WHV, but also contained reads matching to *Herpesviridae, Flaviviridae, Poxviridae* and *Retroviridae* (Figure S9). *O. virginianus* libraries contained *Poxviridae, Retroviridae*, and *Herpesvirida*e reads. Consequently, reads matching to *Poxviridae* were detected in libraries for all deltavirus hosts which were studied here (DrDV-A, DrDV-B, PmacDV, MmonDV, OvirDV), although reads were less abundant than other viral taxa and could not always be decisively ruled out as false positives. Indeed, blastn analysis of poxvirus-like reads, which were originally identified by blastx, revealed that these reads frequently had poor correspondence to *Poxviridae* at the nucleotide level. Although there is no experimental evidence that poxviruses can produce infectious deltavirus particles, this putative ecological association may be worth considering in future studies of mammalian deltaviruses.

## Supplementary Figures

## Supplementary Tables

**Dataset S1 (separate file). Merged Serratus and Pantheria datasets**. Data S1 is an excel file containing merged datasets used to evaluate geographic and taxonomic biases in SRA searches.

**Dataset S2 (separate file). Full delta antigen alignment**. Data S2 is a fasta file which is a trimmed alignment of the amino acid sequence of the full delta antigen used to evaluate relationships between deltavirus representatives from different hosts.

**Dataset S3 (separate file). Vampire bat deltavirus infections**. Data S3 is an excel file containing individual level infection data for vampire bats screened by PCR for DrDV-B along with demographic data used in statistical analyses.

**Dataset S4 (separate file). DrDV-B partial delta antigen alignment**. Data S4 is a fasta file which is a trimmed alignment of the nucleotide sequence of the partial delta antigen fragment used to evaluate relationships between deltaviruses detected in vampire bats from different regions of Peru. A DrDV-B sequence detected in a co-roosting *C. perspicillata* individual is also included in this alignment.

---

**Fig S1.**
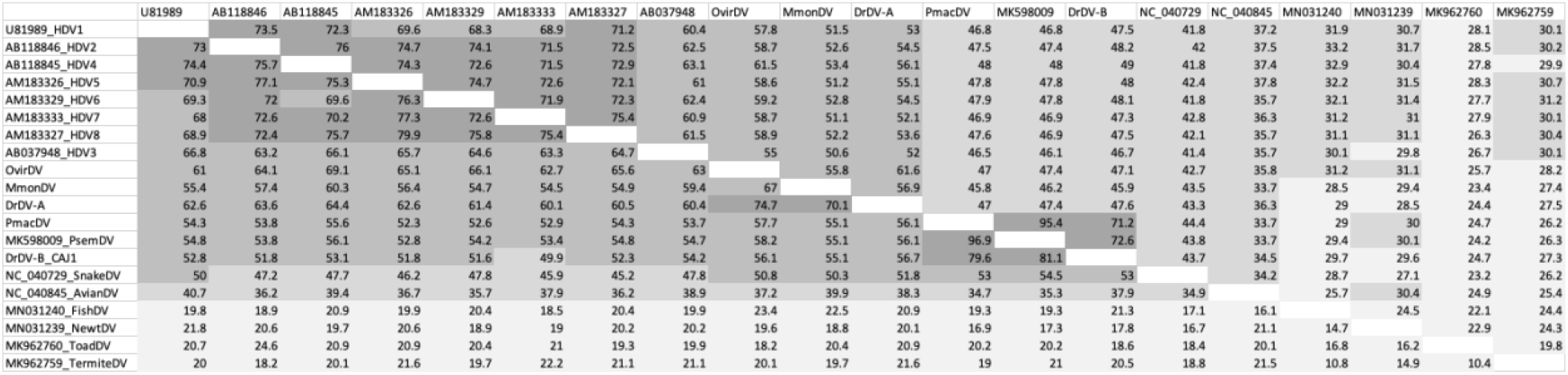
Genetic distances matrices showing representative deltavirus sequences with percent nucleotide identities between genomes (upper triangle) and percent amino acid identities between complete DAg sequences (lower triangle). Darker shading indicates higher percentage identity between two deltaviruses.

**Fig S2.**
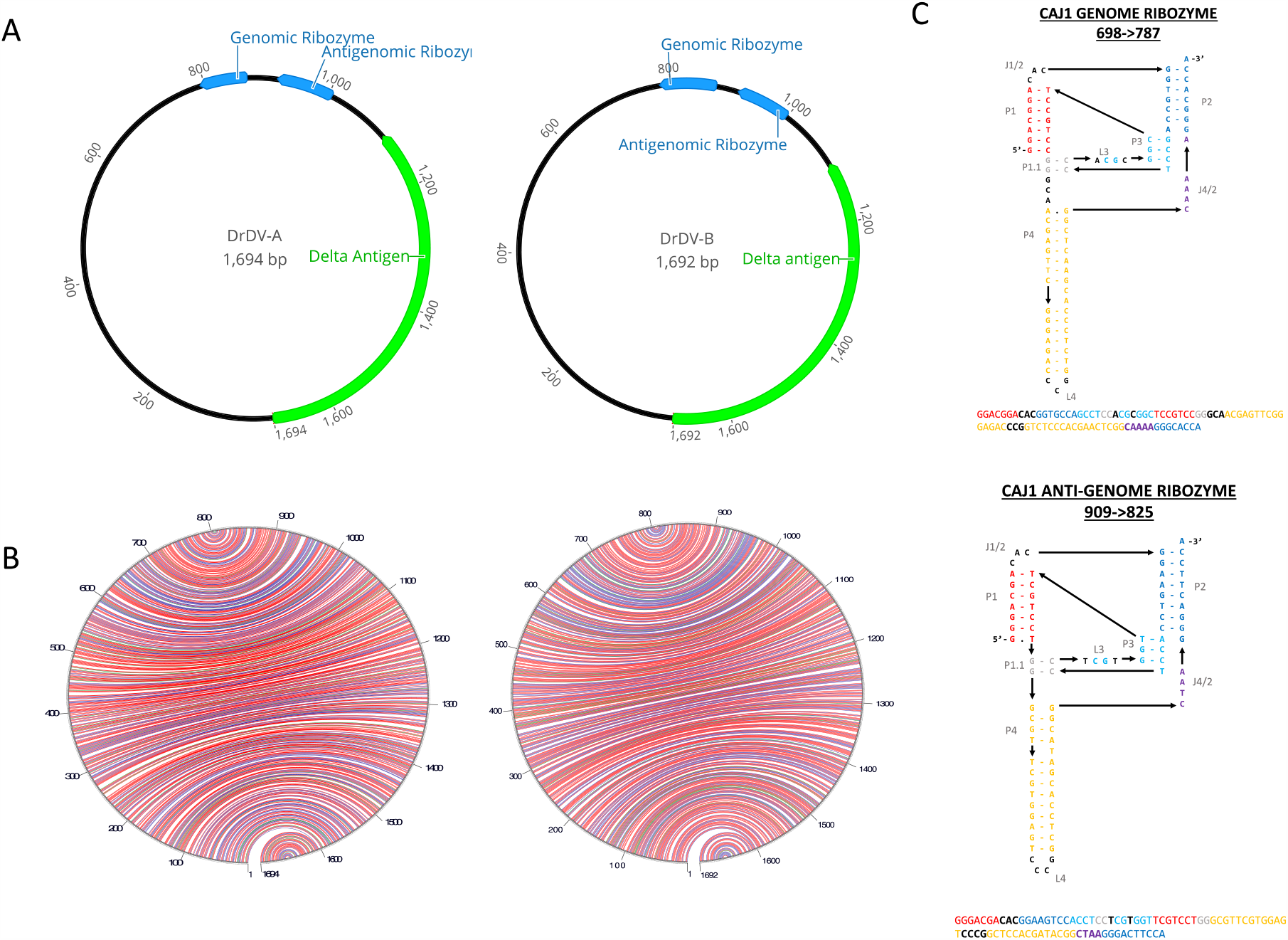
DrDV genomes exhibit characteristics common to deltaviruses. (**A**) The locations of the delta antigen open reading frame (green) and genomic/antigenomic ribozymes (blue) are shown along the circular genomes of DrDV-A and DrDV-B (CAJ1 shown as an example of DrDV-B). (**B**) Intramolecular base pairing for DrDVs depicted as lines connecting points on the circular genome – G-C pairs are red, A-U pairs are blue, G-U pairs are green, other pairs are yellow. (**C**) Genomic and antigenomic ribozyme secondary structures are shown along with genome location for genome CAJ1. Complementary regions are shown in the same color, and structures are depicted in the style of Webb & Luptak to facilitate comparison with ribozymes from previous studies (3, 5, 6). For further comparison, the ribozyme structures presented in (4) are based on a consensus ribozyme sequence created from an alignment of all deltavirus and deltavirus-like ribozymes. Unlike ribozyme sequences in some other deltavirus genomes, we do not observe a shortening of the J1/2 loop in the DrDV genomic ribozyme compared to the anti-genomic ribozyme, with the sequence CAC present in both.

**Fig S3.**
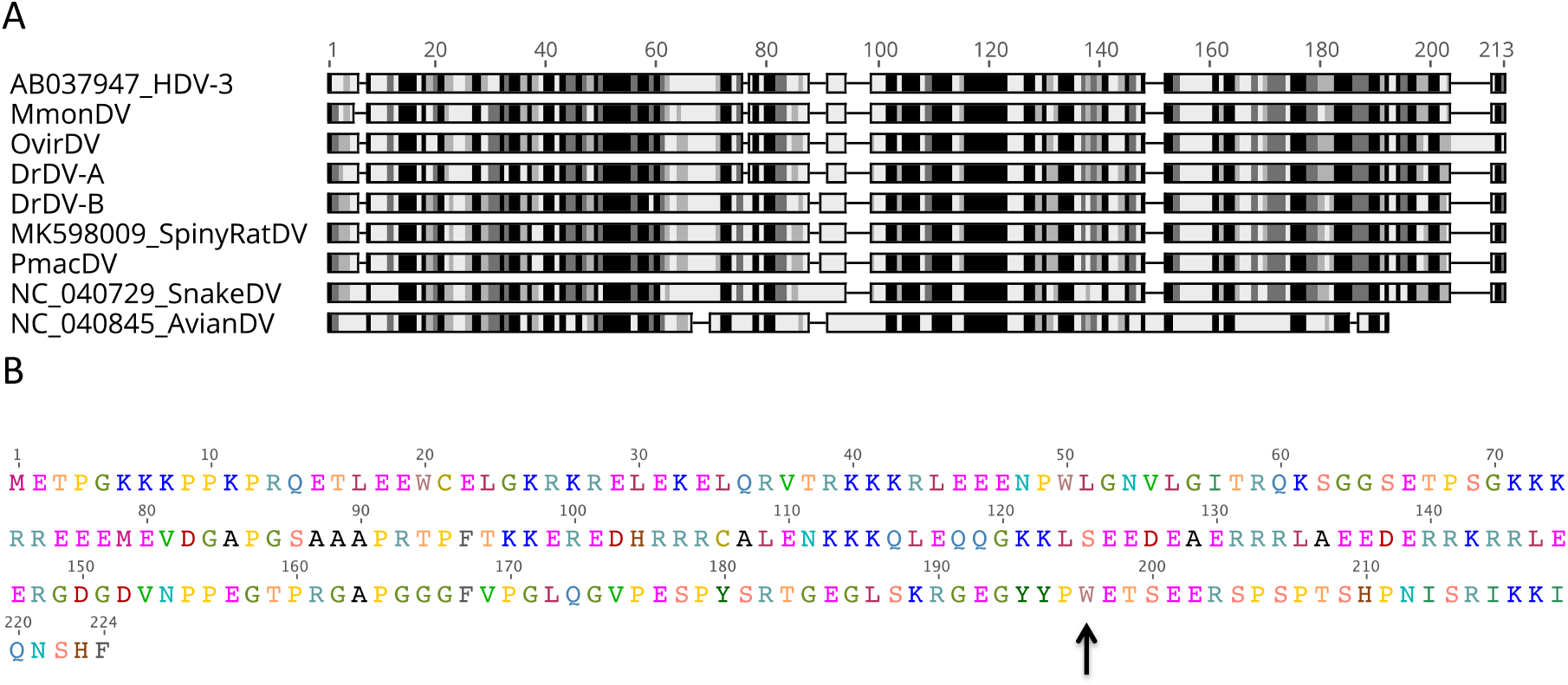
Characterization of DrDV delta antigen proteins. (**A**) Alignment of delta antigen protein sequences for mammalian, snake and avian deltaviruses. Shading indicates level of similarity across all sequences, with regions of highest identity in black. (**B**) Putative sequence of the large DAg for the DrDV-B virus from the site CAJ1. The RNA editing site is marked with a black arrow; UAG has been edited to UGG yielding a tryptophan residue (W).

**Fig S4.**
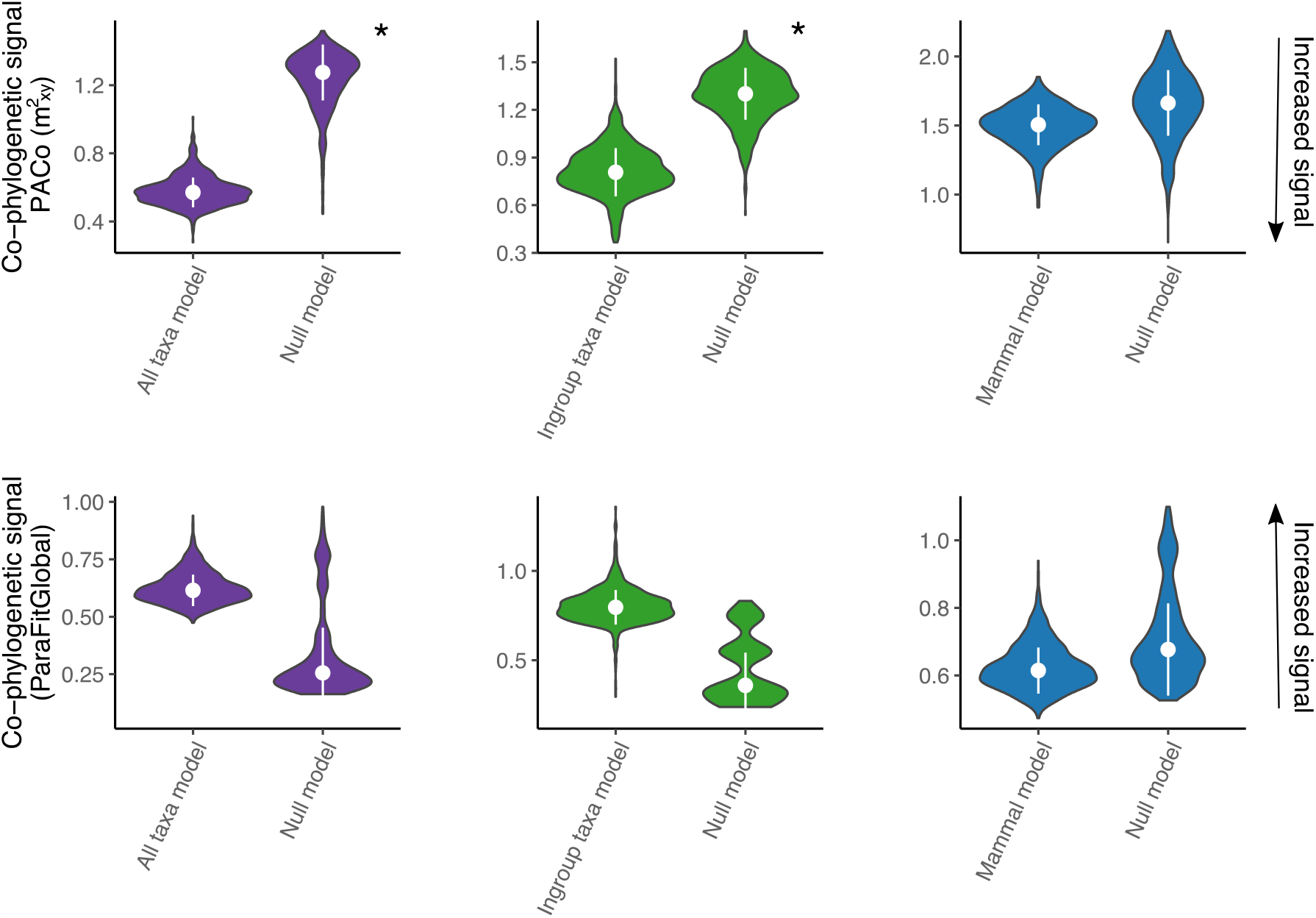
Co-phylogenetic signal in subsets of the deltavirus phylogeny. Violin plots show the degree of dependence of 1,000 phylogenies from the posterior of the StarBeast analysis (Fig. S5) on the host phylogeny relative to null models, with the median and standard deviation. Data subsets are colored as in Fig. 2B (All taxa: purple+green+blue, Ingroup: green+blue, mammals: blue) Distributions are shown for analyses performed using PACo (top row) and ParaFit (bottom row). Asterisks show significant dependence of the virus phylogeny on the host phylogeny (*P* < 0.05). Note that lower values of the empirical model relative to the null model represent increased signal of co-phylogeny in PACo while higher values represent increased signal of co-phylogeny in ParaFit.

**Fig S5.**
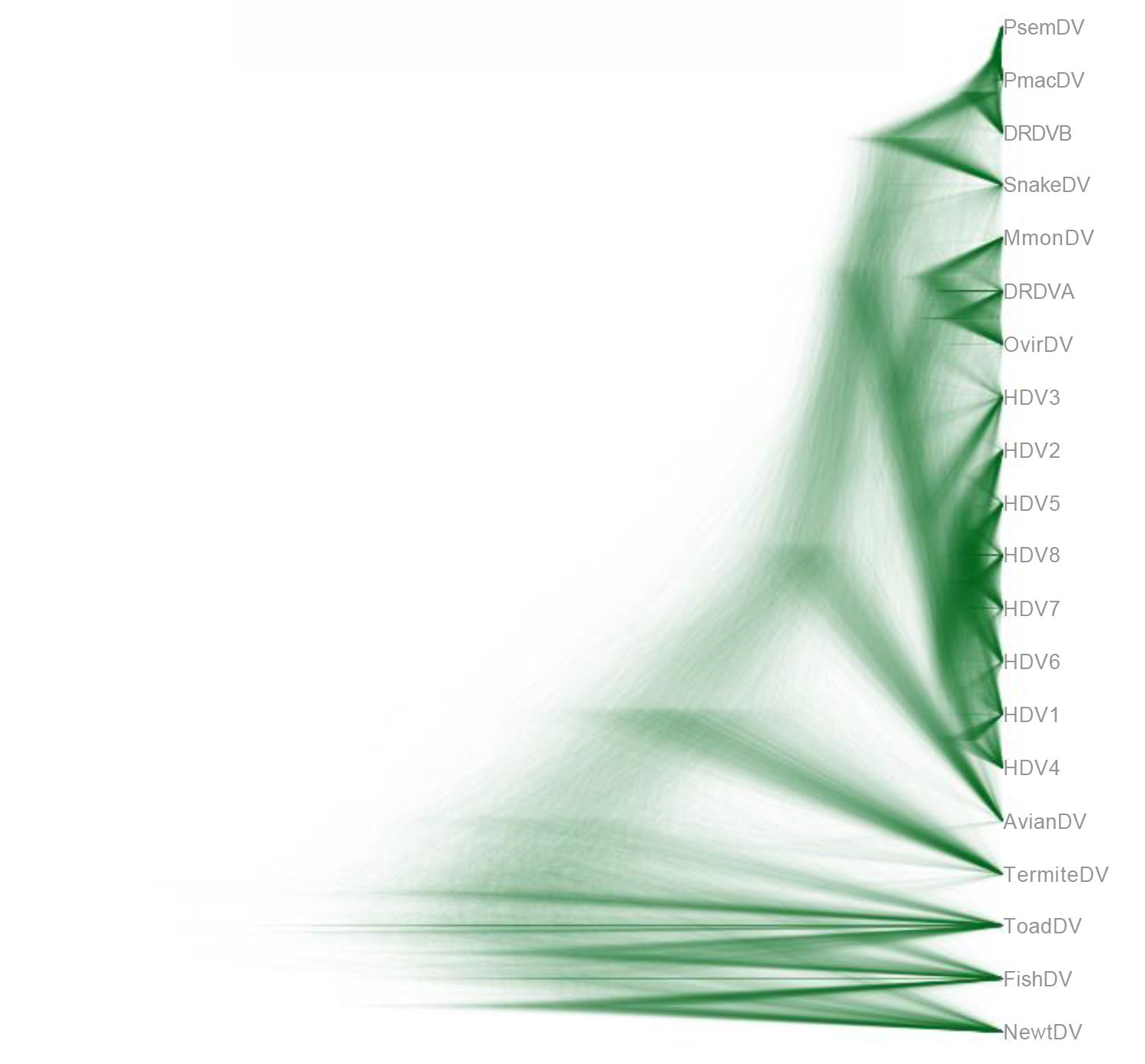
Uncertainty of deep relationships in the deltavirus phylogeny. The DensiTree shows the distribution of 1,000 posterior trees from the StarBeast analysis, highlighting uncertainties in the evolutionary relationships among divergent deltavirus-like taxa.

**Fig S6.**
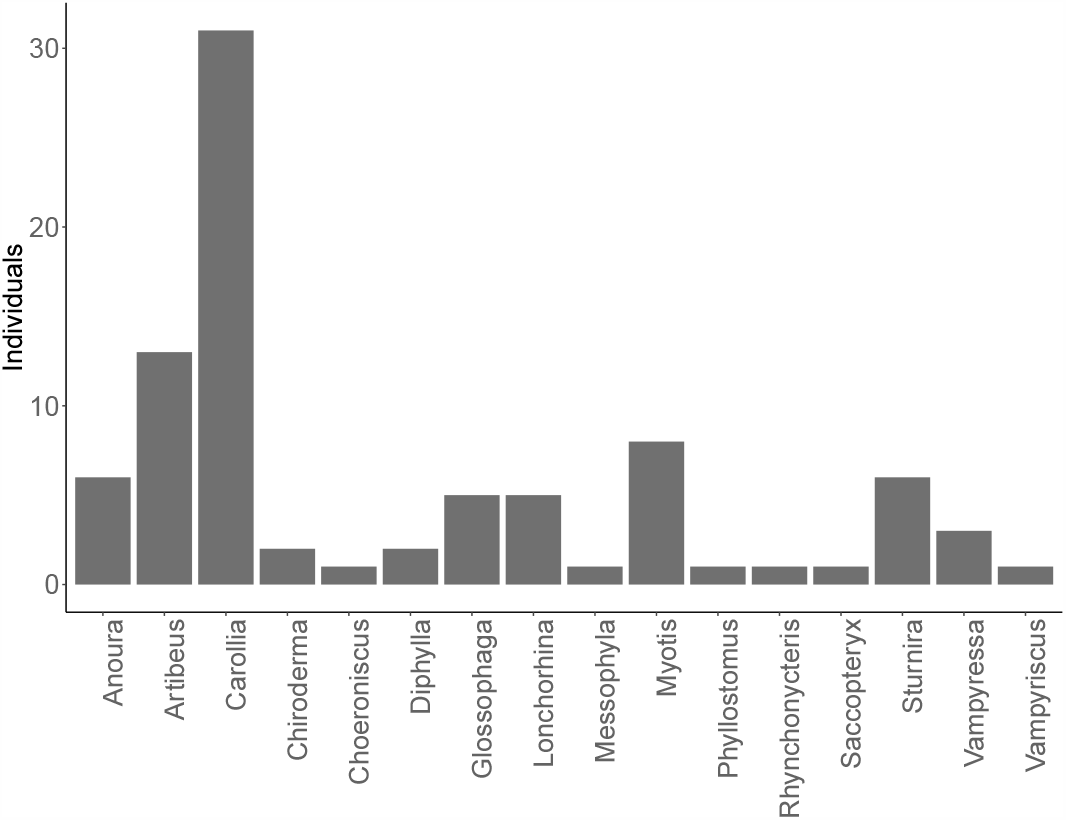
Counts of non-*D. rotundus* bat species saliva swabs individually screened by RT-PCR for DrDV-B. Bars group bats by genus.

**Fig S7.**
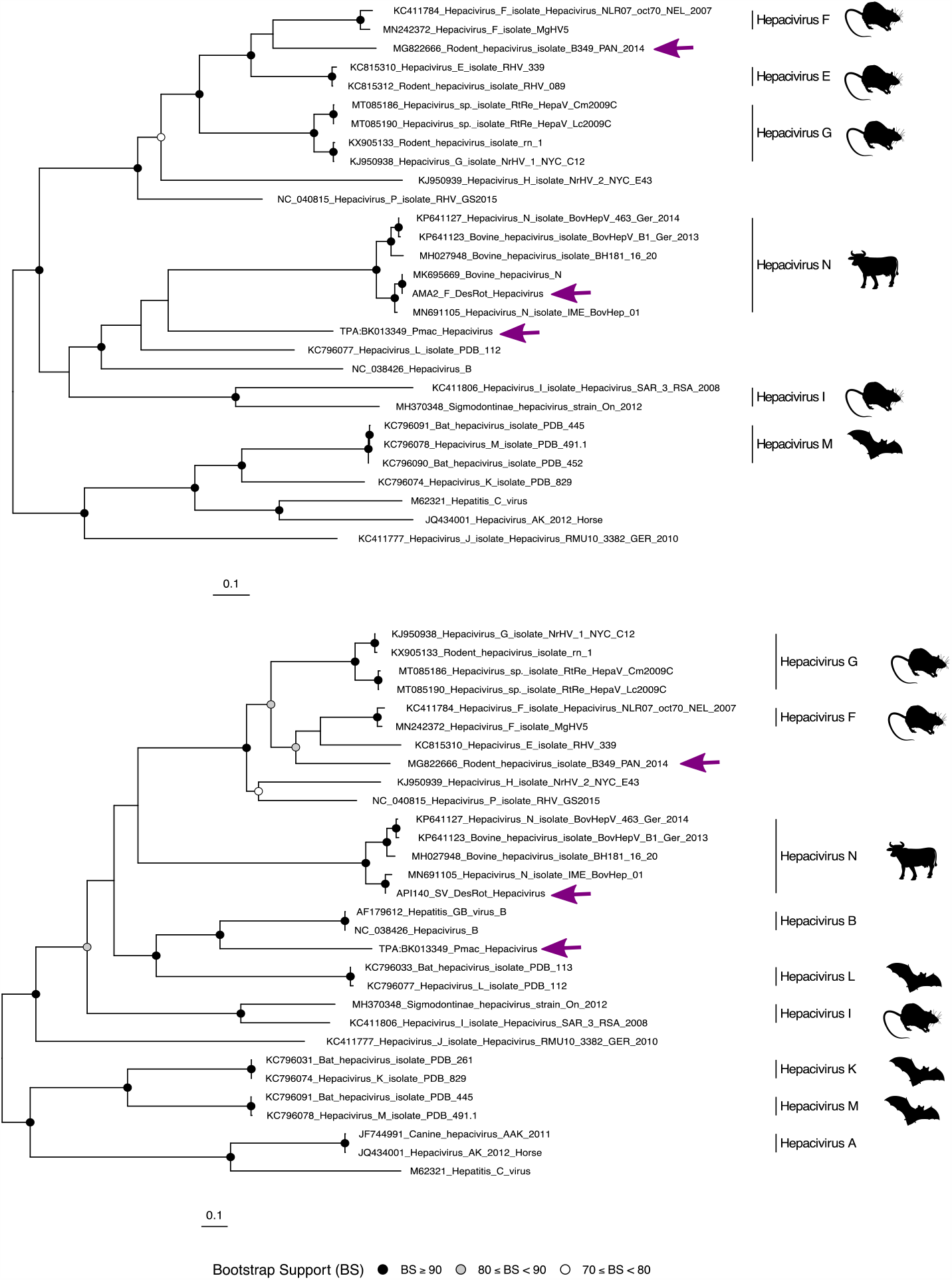
Relationships between hepaciviruses from deltavirus-positive hosts. Phylogenies shown are based on amino acid alignments of the hepacivirus genes NS3 (upper) and NS5B (lower). Hepacivirus species with multiple representatives are denoted with vertical lines. Silhouettes show host associations for key hepacivirus species. Purple arrows indicate hepaciviruses detected in deltavirus-positive hosts (*Peropteryx macrotis, Desmodus rotundus*, and *Proechimys semispinosus*). Maximum likelihood phylogenies with 1,000 bootstrap replicates were generated using IQTree (7) using the best fit models LG+F+I+G4 (NS3) and LG+I+G4 (NS5B) selected by ModelFinder within IQTree 2 (8).

**Fig S8.**
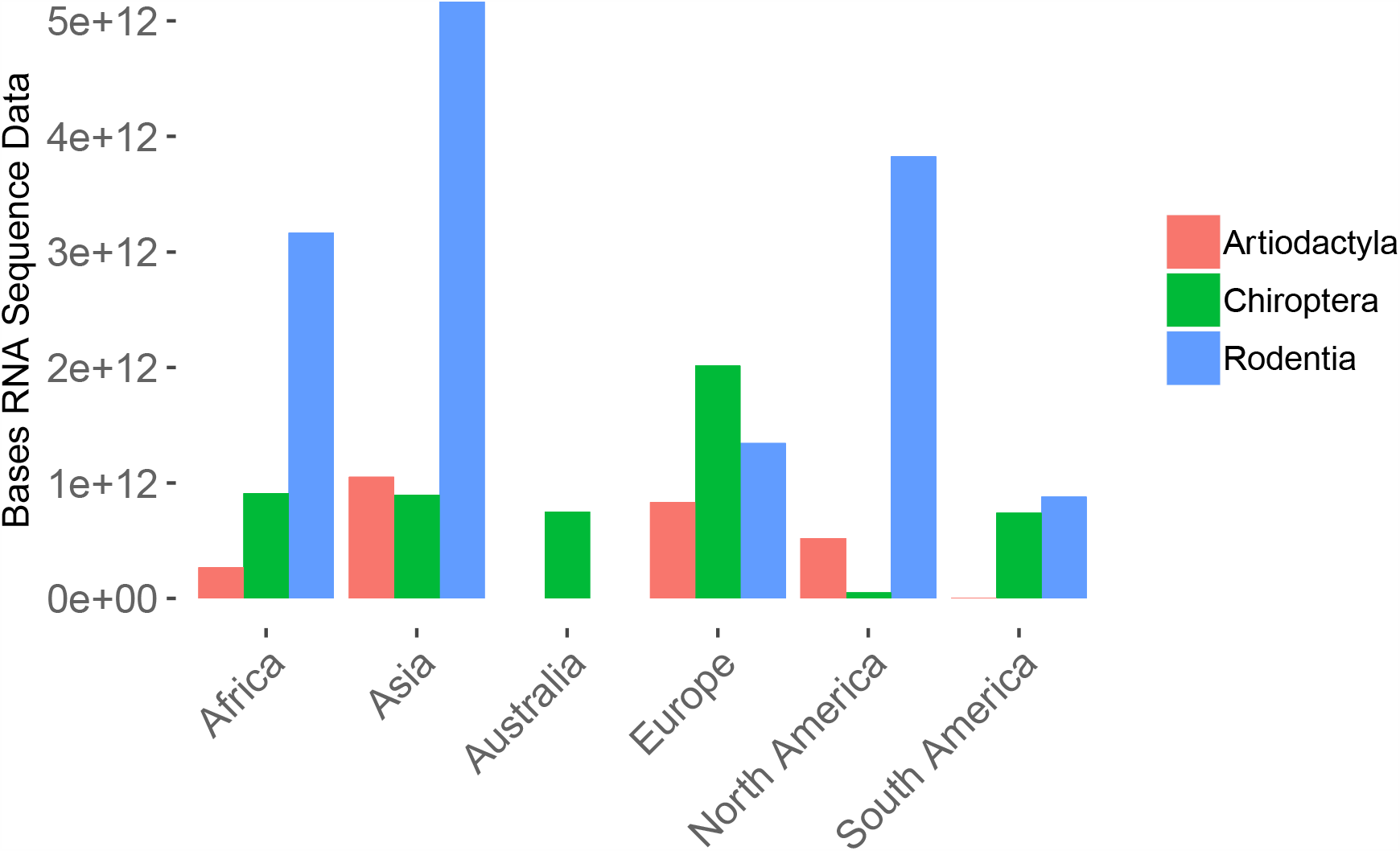
Continent level geographic biases in RNA sequence data examined by Serratus. Bars are colored by mammalian order; data shown are limited to the three orders in which deltaviruses were detected.

**Fig S9.**
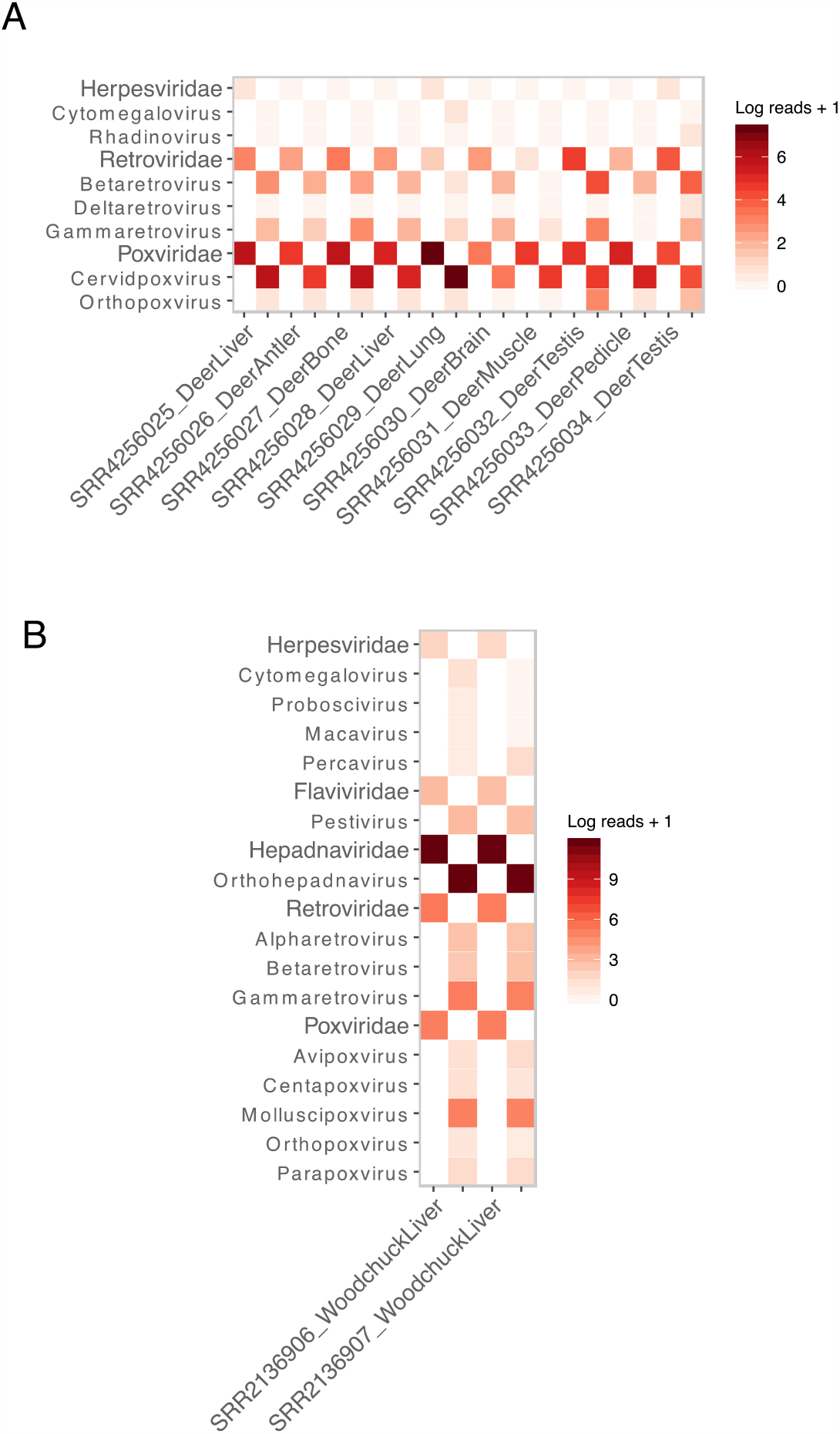
Candidate helper viruses for the OvirDV and MmonDV datasets. Mammal-infecting viral communities are shown for (A) *O. virginianus* libraries sequenced by RNASeq from (9), several of which contained OvirDV and (B) two *M. monax* samples infected with MmonDV from (10). Viral families (in larger font) and genera are shown in adjacent columns for each sample, with families on the left and genera on the right.

**Table S1.** Pooled bat saliva samples from Peru analyzed by metagenomic sequencing.

| Genus | Species | Individuals<br>in pool | Raw reads | Deltavirus<br>contig length<br>(nt) |
| --- | --- | --- | --- | --- |
| *Carollia* | *perspicillata* | 10* | 28,700,978 | N |
| *Glossophaga* | *soricina* | 5 | 24,079,752 | N |
| *Desmodus* | *rotundus* | 10 | 28,946,275 | 921<sup>†</sup> |
| *Diphylla* | *ecaudata* | 2 | 25,023,095 | N |
| *Anoura* | *geoffroyi*<br>*peruana* | 6 | 18,569,505 | N |
| *Artibeus* | *lituratus*<br>*obscurus*<br>*planirostris*<br>*fraterculus* | 10 | 14,966,399 | N |
| *Myotis* | *oxyotus*<br>*unidentified sp* | 8 | 19,934,479 | N |
| *Sturnira* | *erythromos*<br>*unidentified sp* | 6 | 11,348,995 | N |
| *Vampyressa*/<br>*Vampyriscus* | *bidens*<br>*unidentified sp* | 4 | 13,734,389 | N |
| *Rare species* | *Chiroderma trinitatum*<br>*Chiroderma salvini*<br>*Choeroniscus minor*<br>*Rhynchonycteris naso*<br>*Saccopteryx bilineata*<br>*Messophyla macconelli*<br>*Phyllostomus discolor*<br>*Rhinophylla pumilio* | 8 | 16,746,795 | N |

\*Pool included individual CP-1 in which DrDV-B was detected by RT-PCR
<sup>†</sup>Pool was identical to CAJ1 pool where DrDV was initially discovered, confirming the ability to detect deltaviruses when they are known to be present

**Table S2.**
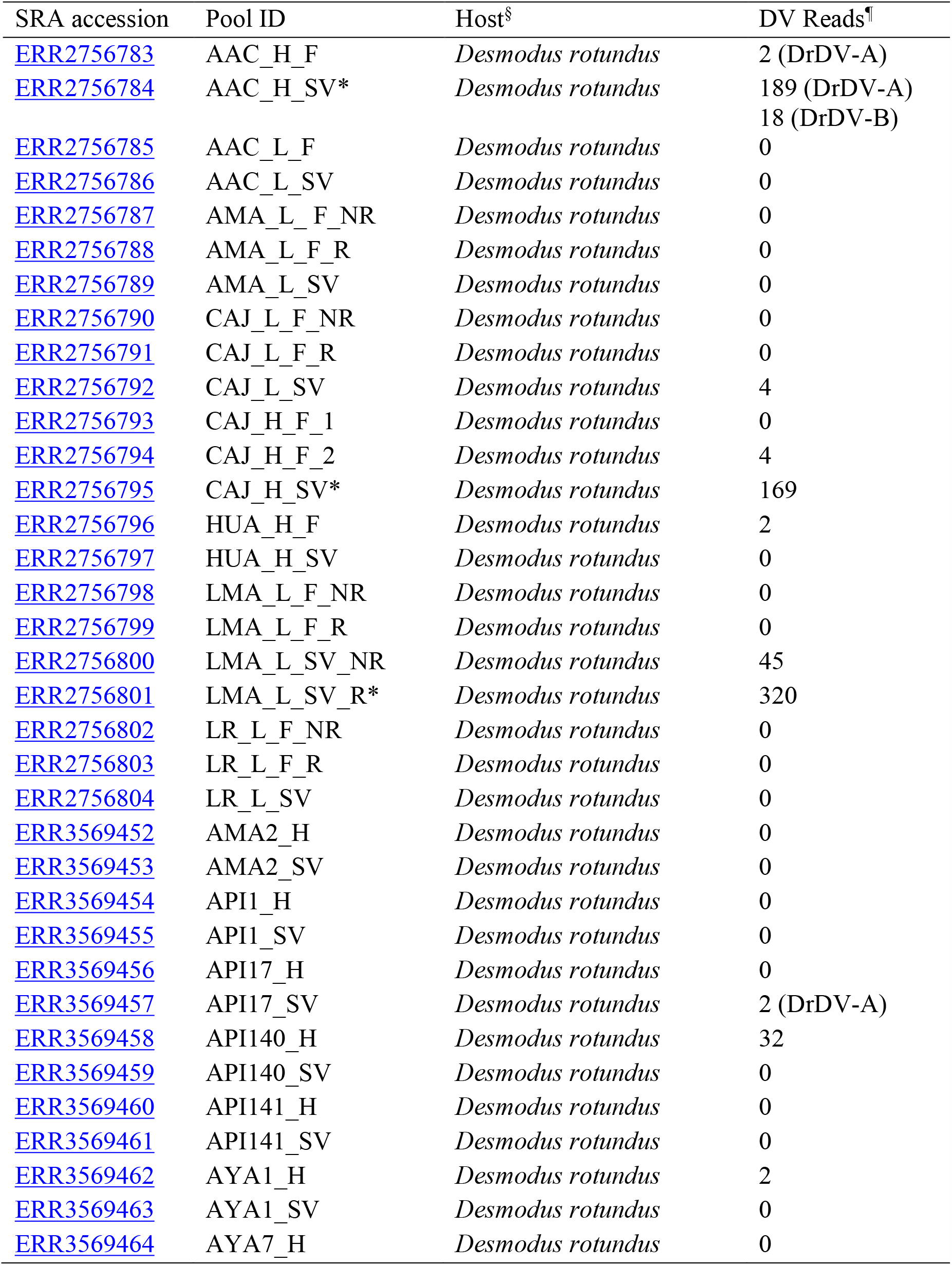
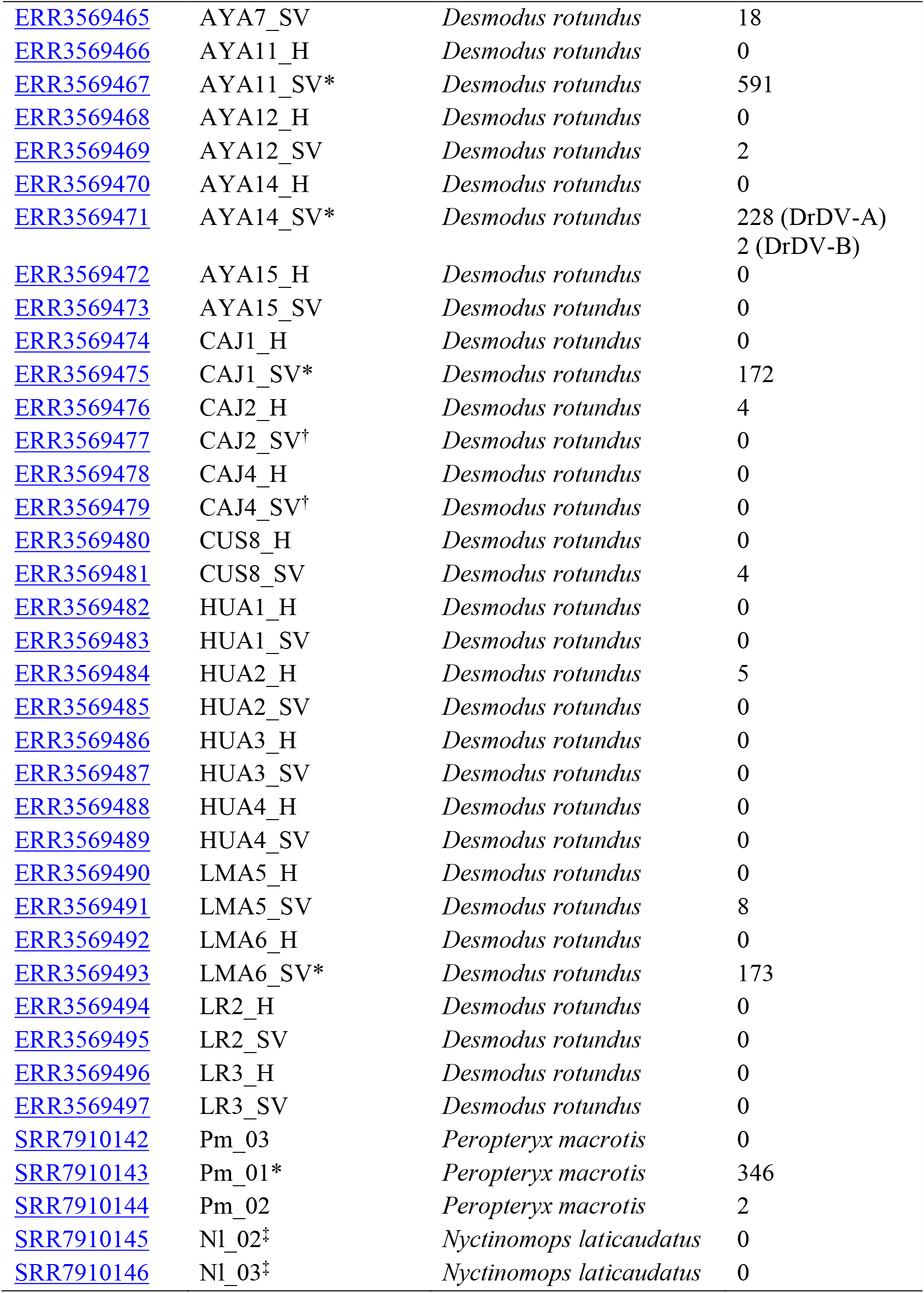
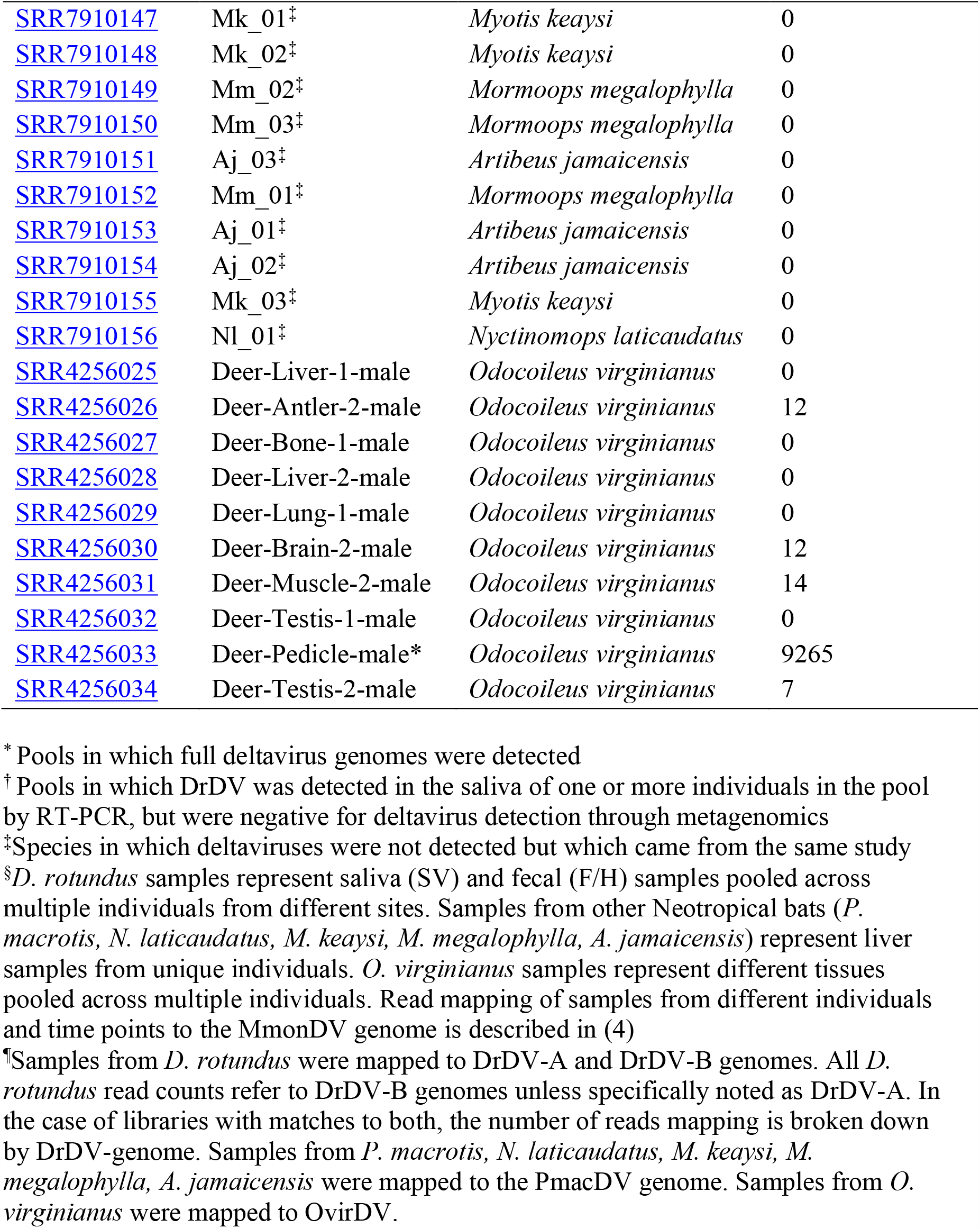
Deltavirus positive cohorts evaluated by mapping reads from related libraries to novel deltavirus genomes.

**Table S3.**
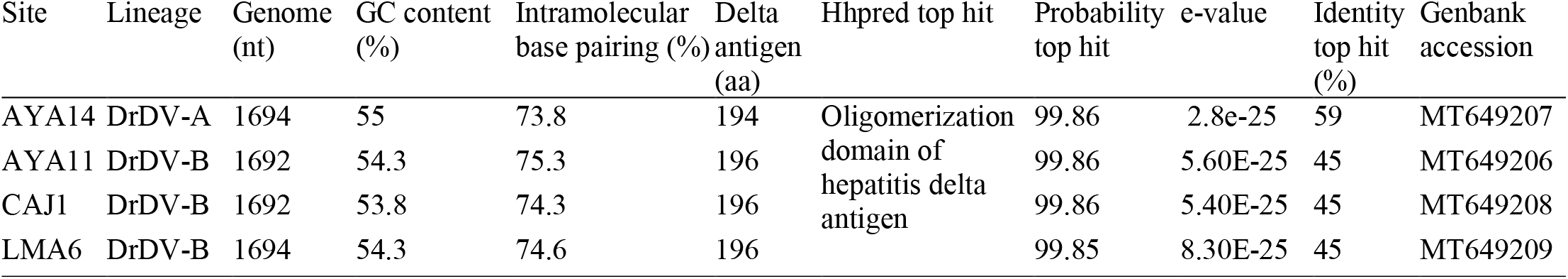
Summary statistics for bat deltavirus genomes and protein domain homology analysis of predicted DrDV small delta antigens from saliva metagenomic pools.

**Table S4.**
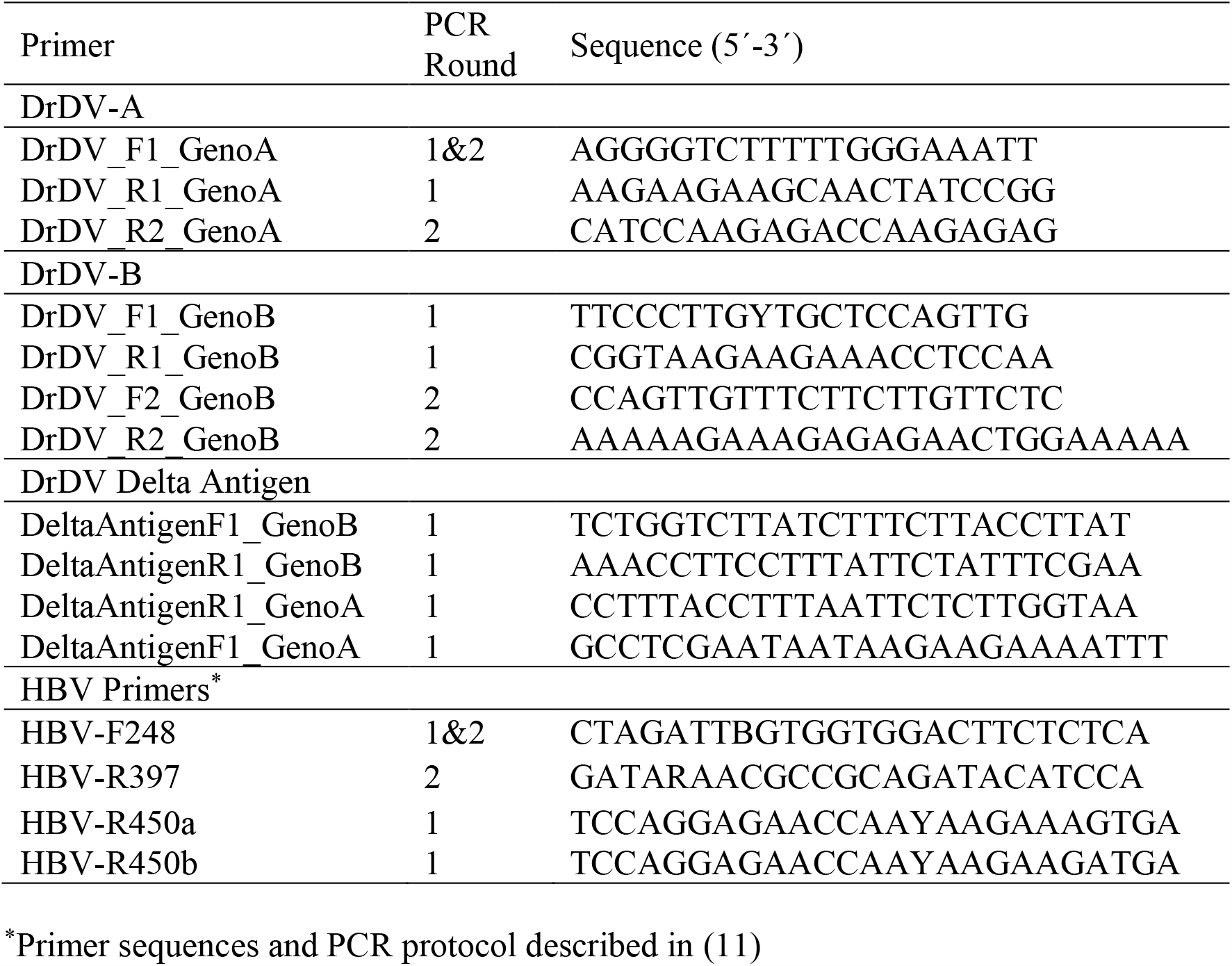
Primers used to screen samples for DrDV by RT-PCR and HBV by PCR.

**Table S5.**
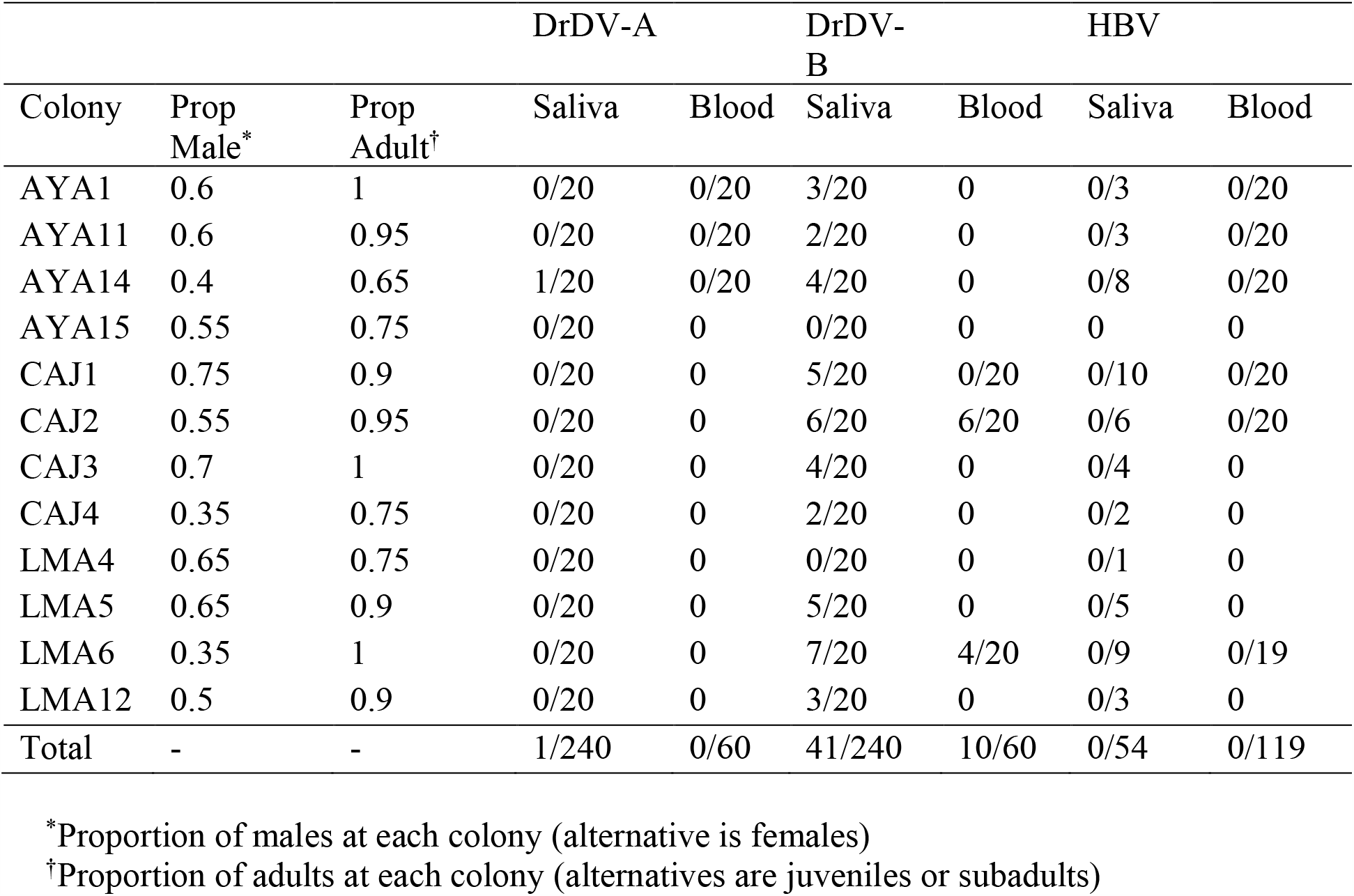
Colony level demographic characteristics and PCR-based screening results of vampire bat blood and saliva for DrDV and HBV.

**Table S6.** Test of association between DrDV-B phylogeny and sample location at the regional (department) and colony level.

| Level | Index | Observed value (95% CI) | Null value (95% CI) | p-value |
| --- | --- | --- | --- | --- |
| Region | AI* | 0.22 (0-0.58) | 2.46 (1.95-2.94) | 0 |
|  | PS† | 4 (3-5) | 17.73 (15.41-19.35) | 0 |
|  | MC‡ (LMA) | 9.29 (5-14) | 1.97 (1.41-2.98) | 0.001 |
|  | MC‡ (CAJ) | 11.24 (9-19) | 2.81 (2.12-3.94) | 0.001 |
|  | MC‡ (AYA) | 2.67 (1-5) | 1.26 (1-1.96) | 0.02 |
| Colony | AI* | 2.23 (1.69-2.76) | 3.63 (3.21-3.92) | 0 |
|  | PS† | 19.19 (18-20) | 29.23 (27.45-30.81) | 0 |
|  | MC‡ (LMA6) | 4.98 (5-5) | 1.18 (1-1.94) | 0.001 |
|  | MC‡ (LMA5) | 2.52 (1-3) | 1.13 (1-1.43) | 0.002 |
|  | MC‡ (CAJ2) | 3.21 (2-6) | 1.57 (1.14-2.39) | 0.01 |
|  | MC‡ (CAJ1) | 1 (1-1) | 1.13 (1-1.53) | 1 |
|  | MC‡ (AYA1) | 1.09 (1-2) | 1.01 (1-1.05) | 1 |
|  | MC‡ (CAJ3) | 1.01 (1-1) | 1.07 (1-1.32) | 1 |
|  | MC‡ (AYA11) | 1.03 (1-1) | 1.01 (1-1.05) | 1 |
|  | MC‡ (AYA14) | 1.05 (1-2) | 1.04 (1-1.15) | 1 |
|  | MC‡ (CAJ4) | 1.16 (1-2) | 1.01 (1-1.05) | 1 |
|  | MC‡ (LMA12) | 1 (1-1) | 1.04 (1-1.15) | 1 |

\* Association Index
† Parsimony Score
‡ Monophyletic Clade size

